# Self-growing protocell models in aqueous two-phase system induced by internal DNA replication reaction

**DOI:** 10.1101/2024.07.01.599542

**Authors:** Yoshihiro Minagawa, Moe Yabuta, Masayuki Su’etsugu, Hiroyuki Noji

## Abstract

The bottom-up reconstitution of self-growing artificial cells is a critical milestone toward realizing autonomy and evolvability. However, building artificial cells that exhibit self-growth coupled with internal replication of gene-encoding DNA has not been achieved yet. Here, we report self-growing artificial cell models based on dextran-rich droplets in an aqueous two-phase system of poly(ethylene glycol) (PEG) and dextran (DEX). Motivated by the finding that DNA induces the generation of DEX-rich droplets, we integrated DNA amplification system with DEX-rich droplets, which exhibited active self-growth. We implemented the protocells with cell-free transcription-translation (TXTL) systems coupled with DNA amplification/replication, which also showed active self-growth. We also observed self-growth activity of protocells carrying a single copy of DNA. Considering the simplicities in terms of the chemical composition and the mechanism, these results underscore the potential of DEX droplets as a foundational platform for engineering protocells, giving implications for the emergence of protocells under prebiotic conditions.

## Introduction

The bottom-up reconstitution of autonomous artificial cell systems from natural and/or synthetic components is one of the grand challenges in synthetic biology. Artificial cell models with essential features of living cells, such as replication of genetic materials, gene expression, information/energy transductions, and vesicle fission/division, have been developed to elucidate the working mechanism of molecular systems (1–4). The bottom-up approach of autonomous artificial cell systems also provides implications for possible scenarios on how protocells have emerged in prebiotic conditions (5). From the engineering point of view, the open nature of artificial cell systems holds a great promise for technological innovations in the rapid prototyping or high-throughput screening of functional biomolecules/biosystems that would be applicable in various fields such as bioanalysis, medicine, and material science (6–10).

Autonomous artificial cell models should contain a genetic information-carrying molecule and a set of catalyst molecules that replicate genetic molecules and produce the catalyst molecules themselves. In addition, genetic molecules and catalyst molecules should be micro-compartmentalized in a microreactor as a ‘cell’ to define the identity of the cell and protect itself from inevitably emerging parasites (11). Considerable progress has been made in the reconstitutions of individual subcellular systems such as DNA/RNA amplification/replication, coupled transcription and translation, lipid synthesis, and lipid vesicle division machinery (12–18). However, there are still large gaps between natural and artificial cells in terms of the orchestration of multiple functions to realize autonomy. In particular, the growth and division of micro-compartmentalizing vesicles or reactors coupled with the replication of genetic materials remain as a grand challenge.

Membrane vesicles composed of fatty acids or phospholipids are the gold standard for the micro-compartmentalization in artificial cell research. Dynamic morphological changes in vesicles, such as growth, fission, and division, were observed when fatty acids or lipids were externally or enzymatically supplied or when the excluded volume effect of DNA was applied to the lumen of the vesicle (14,19–23). There are also studies on the enhancement of vesicle budding or division after in-vesicle DNA amplification using PCR, accompanied by the generation of vesicle-forming molecules in the presence of the catalyst (24–26). However, self-growing vesicles coupled with or driven by nucleic acid amplification without external controls have not yet been developed.

Another class of micro-compartmentalization methods that have recently attracted growing attention is liquid–liquid phase separation (LLPS) (2,4,27). Microdroplets formed via LLPS have no physical boundary between droplets and external continuous phase, thus enabling solute exchange with the continuous phase and allowing a continuous supply of substrate molecules, such as nucleic acids and amino acids, from the external solution. In addition, some types of LLPS droplets can enrich the biopolymers inside them. Taking these advantageous features, various studies were reported to attempt to construct artificial cell models. In particular, coacervates were often used (4,28). Several studies have reported that the growth of coacervate droplets by producing constituting molecules of coacervates (29–31).

In contrast, aqueous two-phase system (ATPS) has two solution constituents, in which polymers become immiscible at over threshold concentrations. The ATPS of dextran (DEX) and poly(ethylene glycol) (PEG) is one of the best characterized. It is currently attracting great interest because the DEX-rich phase preferentially enriches nucleic acid polymers, such as DNA and RNA, as well as some types of folded proteins (32–36). The *in vitro* transcription and translation (TXTL) within the DEX-rich droplets in PEG/DEX ATPS have also been reported (37). Thus, LLPS systems have been recognized as an alternative to liposomes for the generation of artificial cell models.

In this study, we first investigated the interplay between PEG/DEX ATPS and DNA. Then, we constructed an ATPS-based artificial cell reactor that exhibits autonomous growth of the DEX-droplet reactor coupled with internal DNA replication or RNA production. We also tested the artificial cell reactor model of DEX-rich droplets that undergoes TXTL coupled DNA amplification/replication to drive the active growth of the cell reactor. We also observed self-growth activity of protocells carrying a single copy of DNA. Our findings can pave the development of artificial cell models with more autonomy.

## Results

We used DEX (MW: 550 kDa) and PEG (MW: 35 kDa) in this study. The composition of the PEG/DEX mixture is presented as a coordinate for simplicity; for example, (*4.0, 4.0*) means a mixture of 4.0% w/w DEX and 4% w/w PEG. First, we quantitatively measured the partitioning ratio of dsDNA in the DEX-rich phase of the PEG/DEX ATPS (*4.0*, *4.0*) (Fig. 1a) from the fluorescence signal of SYBR Gold according to calibration curves (Supplemental Fig. 1). Linear DNAs with lengths of 10, 100, plasmid DNAs 4 × 10^3^, and 205 × 10^3^ bp were tested. The partitioning ratio of DNA in the DEX-rich phase increased with DNA length (Fig. 1b). 4 kbp and 205 kbp DNAs were almost exclusively enriched in the DEX phase, whereas the 10 bp DNA was evenly partitioned in the DEX-rich and PEG-rich phases.

**Fig. 1.**
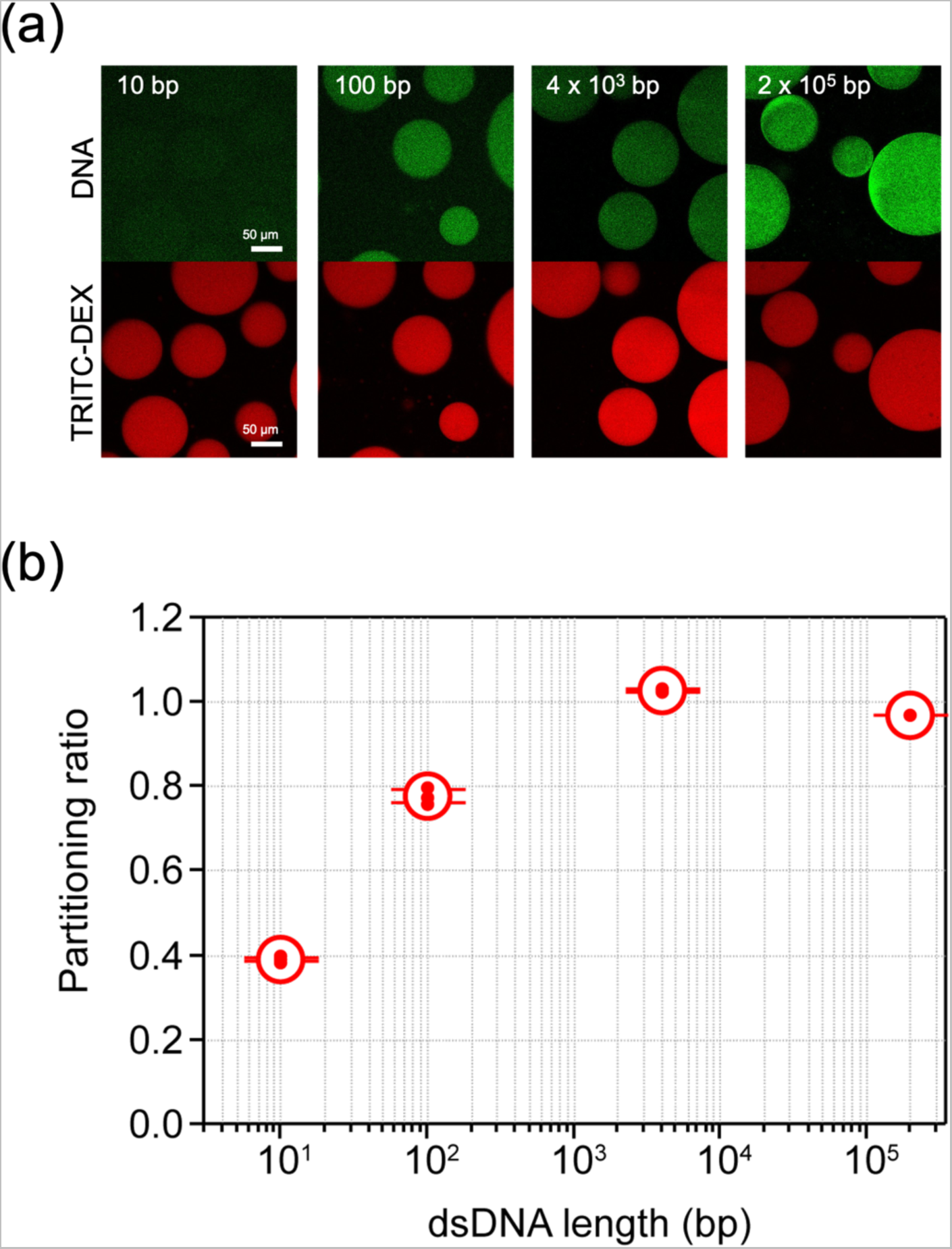
DNA partitioning in PEG/DEX ATPS. (a) Confocal fluorescence images of DNA stained with SYBR Gold in (*4.0, 4.0*) PEG/DEX mix (i.e., 4.0 w/w% PEG and 4.0 w/w% DEX). Concentrations are expressed as coordinates. (b) Partitioning ratio of DNA between the DEX-rich and PEG-rich phases. 10 bp, 100 bp, 4 kbp, and 205 kbp double-stranded DNAs were tested. Error bars represent standard deviation of three experiments.

Then, we examined whether DNA induces phase separation of the PEG/DEX system by comparing phase transition of PEG/DEX systems with or without DNA of 4 or 205 kbp at 950 ng/µL. We employed titration experiment (38), where PEG/DEX ATPS solution was iteratively diluted by pure water to determine the point at which ATPS turned to a miscible solution. Fig. 2a shows a representative result of titration experiment starting from (*2.3, 2.3*). Upon dilution, PEG/DEX ATPS without DNA became miscible at (*2.12, 2.12*), whereas the solution with 4 kbp DNA became a single phase at (*1.98, 1.98*). The solution with 205 kbp DNA underwent phase transition at (*1.85, 1.85*). These results show that DNA stabilizes PEG/DEX ATPS. The titration experiment was repeated three times for each DNA length to determine the probability of the phase transition (Fig. 2b) because phase transition stochastically occurred around the threshold concentration. This is attributable to the subtle differences in experimental conditions and/or the intrinsic stochasticity of PEG/DEX phase transition. We determined the point for a 50% probability of phase transition by fitting the data points with a hyperbolic tangent function. We conducted titration experiments starting from six different compositions (Supplemental Fig. 2) to draw the binodal curves in the phase diagram (Fig. 2c), which clearly showed that DNA stabilizes PEG/DEX ATPS irrespective of the initial conditions of titration experiments. In particular, 205 kbp DNA showed a significant effect on the stabilization of phase separation.

**Fig. 2.**
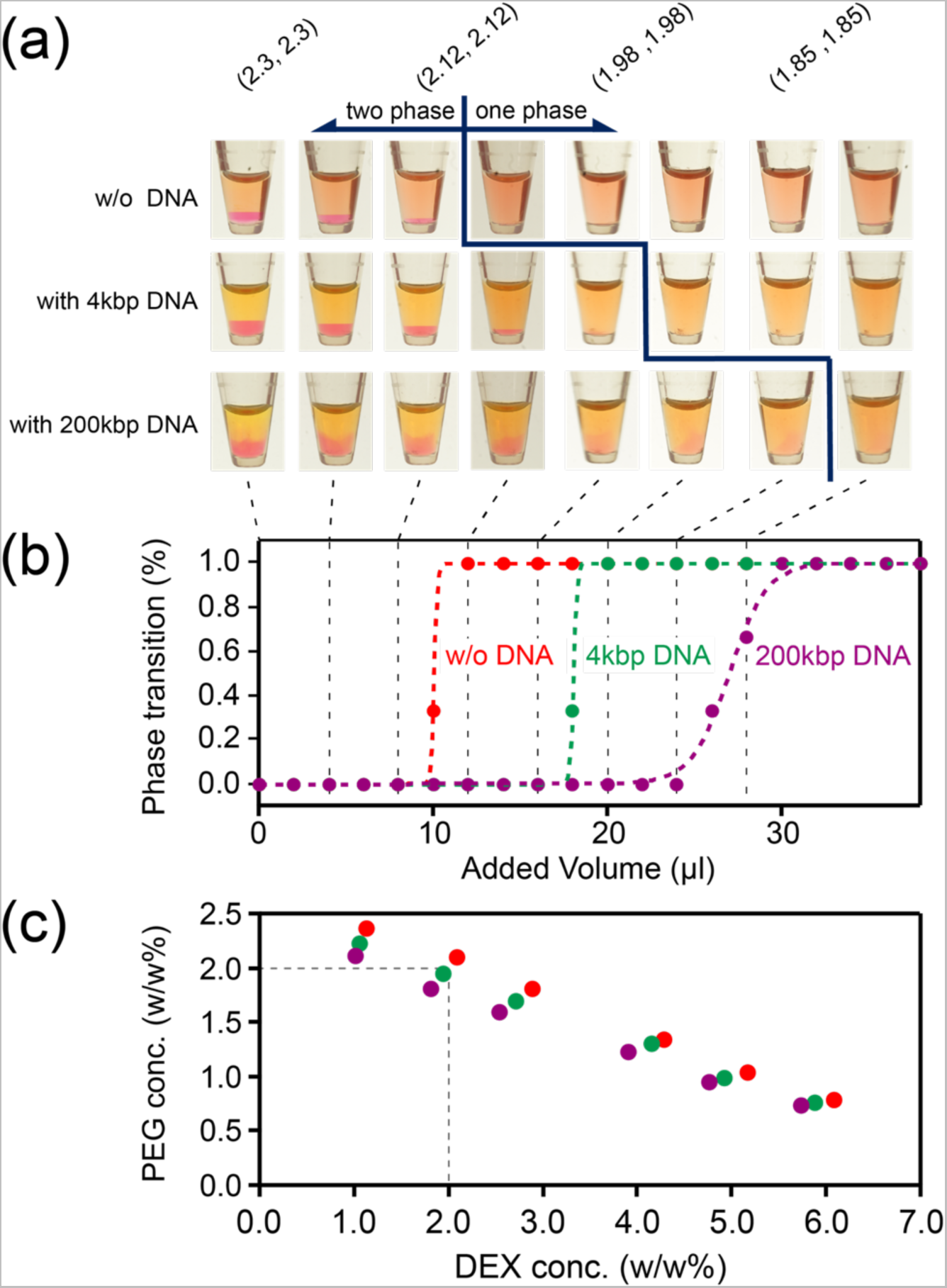
Titrimetry of PEG/DEX ATPS with or without DNA. (a) Photos of the titration experiment of PEG/DEX ATPS with DNA. PEG/DEX mix of (*2.3, 2.3*) was serially diluted by adding small amount of pure water (for details, see Methods). The upper brackets indicate PEG/DEX compositions during titration. The PEG-rich layer (top layer in a tube) and the DEX-rich layer (bottom layer in a tube) were stained with TRITC-DEX and FITC-PEG, respectively. (b) Probabilities of phase transition determined from three experiments. The data points were fitted on hyperbolic tangent functions to determine the critical concentrations (Supplemental Fig. 2). (c) The critical concentrations for the phase transition in the absence or presence of DNA: red, without DNA; green, with 4 kbp DNA; purple, with 205 kbp DNA. The cross point of dashed lines indicates (*2.0, 2.0*) which lies between the critical concentrations with 4 kbp DNA and without DNA. This condition was employed in the following experiments for ATPS induction.

The results of the titration experiments suggest the possibility that DNA amplification can induce the formation of DEX-rich droplets from miscible PEG/DEX solutions. To test this idea, we microscopically observed the PEG/DEX solution in the presence of a DNA amplification reaction mix. The initial PEG/DEX concentrations were set at (*2.00, 2.00*) which is between the binodal curves w/ 4 kbp DNA or w/o DNA as indicated by the crosspoint of dash lines in Fig. 2c: (*2.09, 2.09*) for w/o DNA and (*1.92, 1.92*) for w/ 4 kbp DNA. The reaction mixture of rolling circle amplification (RCA) with EquiPhi29 DNA polymerase was added to the PEG/DEX mix with circular template DNA of 205 kbp at 8 pM (1 ng/μL) (Fig. 3a). In this study, EquiPhi29 DNA polymerase (Equi*ϕ*29DNAP) (39), an activity-enhanced mutant of Phi29 DNA polymerase was used in RCA throughout the experiments. Note that the initial template DNA concentration was sufficiently low to prevent spontaneous phase separation. The PEG/DEX mixture was introduced into a flow cell chamber and observed under a fluorescence microscope. After incubation at 33 °C for 16 h, many DEX-rich droplets with the diameter up to 40 μm appeared (Fig. 3b). Hereafter, DEX-rich droplet is referred to as DEX droplet for simplicity. All DEX droplets enriched DNA inside. When PEG or DEX was absent in the solution, droplets were not observed (Supplemental Fig. 3). DEX droplet generation was also observed when 12 kbp DNA was used as template DNA. In both experiments, time-lapse images showed that DEX droplets appeared and actively coalesced with other droplets (Supplemental Video 1). These observations demonstrated that DNA amplification induces the phase separation of PEG/DEX solution and the growth of DEX droplets.

**Fig. 3.**
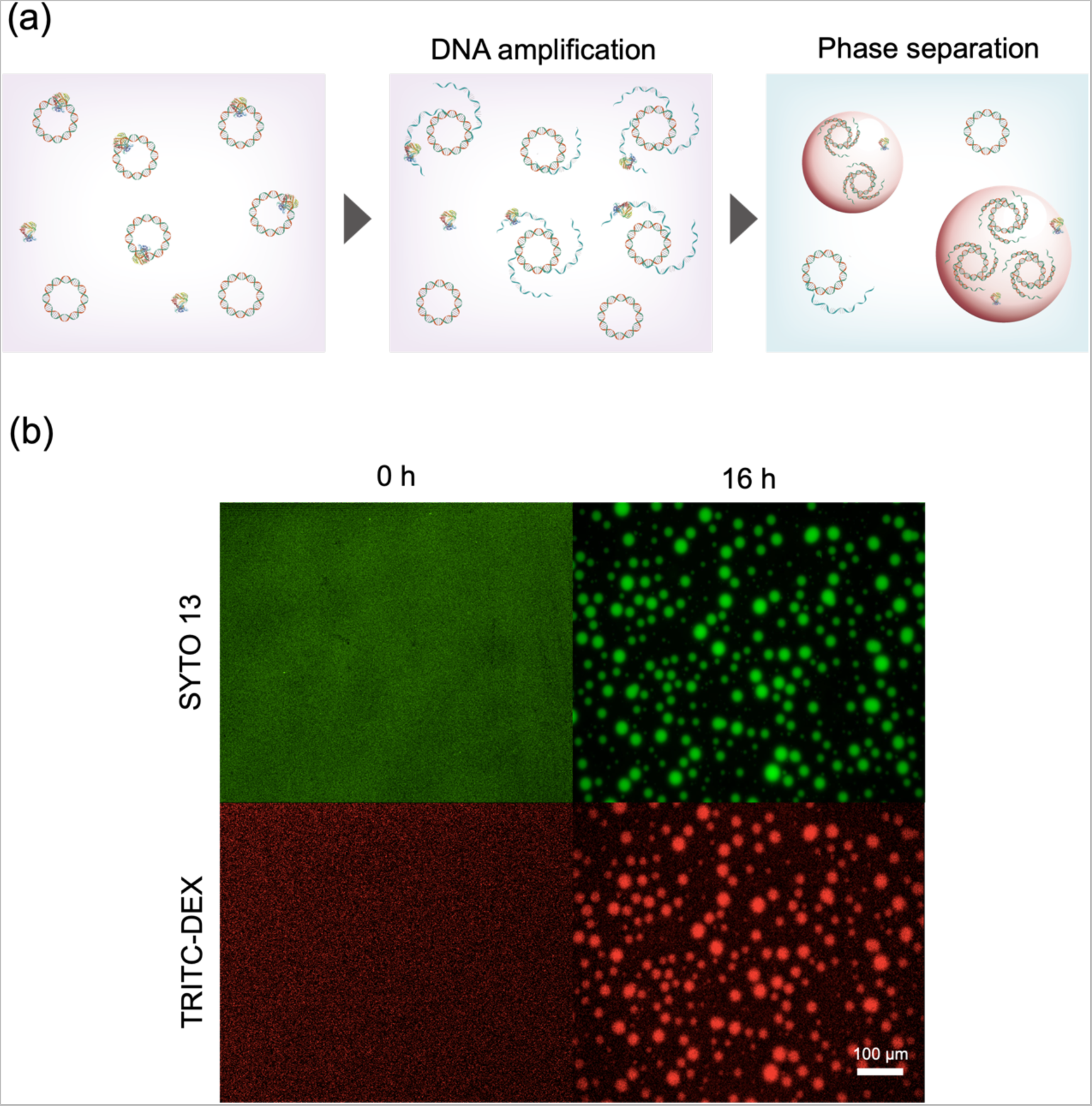
Phase separation induced by RCA. (a) Experimental diagram of phase separation induced by RCA. RCA was conducted with Equi*ϕ*29DNAP in miscible PEG/DEX mix (2.00, 2.00). (b) Fluorescence images show the reaction at the start (0 h, left) and at 16 h (right). DNA and DEX were stained with SYTO13 (green, top) and TRITC-DEX (red, bottom).

For quantitative analysis on the active growth of DEX droplets, we conducted time-lapse observation of DEX droplets (Fig. 4a). DEX droplets in PEG/DEX ATPS solution were prepared and preloaded with RCA reaction mixture containing template 4.2 kbp circular DNA. After introduced into a flow cell, DEX droplets spontaneously settled out due to higher density compared to PEG-rich solution, and stably attached to the coverslip, allowing for time-lapse imaging. While the initial size of DEX droplet differed in each preparation, the diameter ranged typically from 4 to 15 µm with the volume from 30 to 1800 fL (Supplemental Fig. 4a). Upon the incubation at 30 °C, the droplets showed active growth (Fig. 4b and Supplemental Video 2). Droplets showed clear volume expansion; most of droplets increased the volume 2-fold and some showed over 7-fold expansion. After 200 min, most of droplets reached a plateau size pausing the growth, probably due to the complexity nature of RCA; RCA yields dendritic DNA clusters as the reaction product, hampering further DNA amplification. We observed moderate tendency that smaller droplets were more active for volume expansion (Supplemental Fig. 4b). Note that small aggregates were formed on the surface of DEX droplets for unknown reasons in the presence of fluorescently labelled DEX (TRITC-DEX) at 0.05 w/w%. Therefore, TRITC-DEX was added at 0.01 w/w% in the following experiments.

**Fig. 4.**
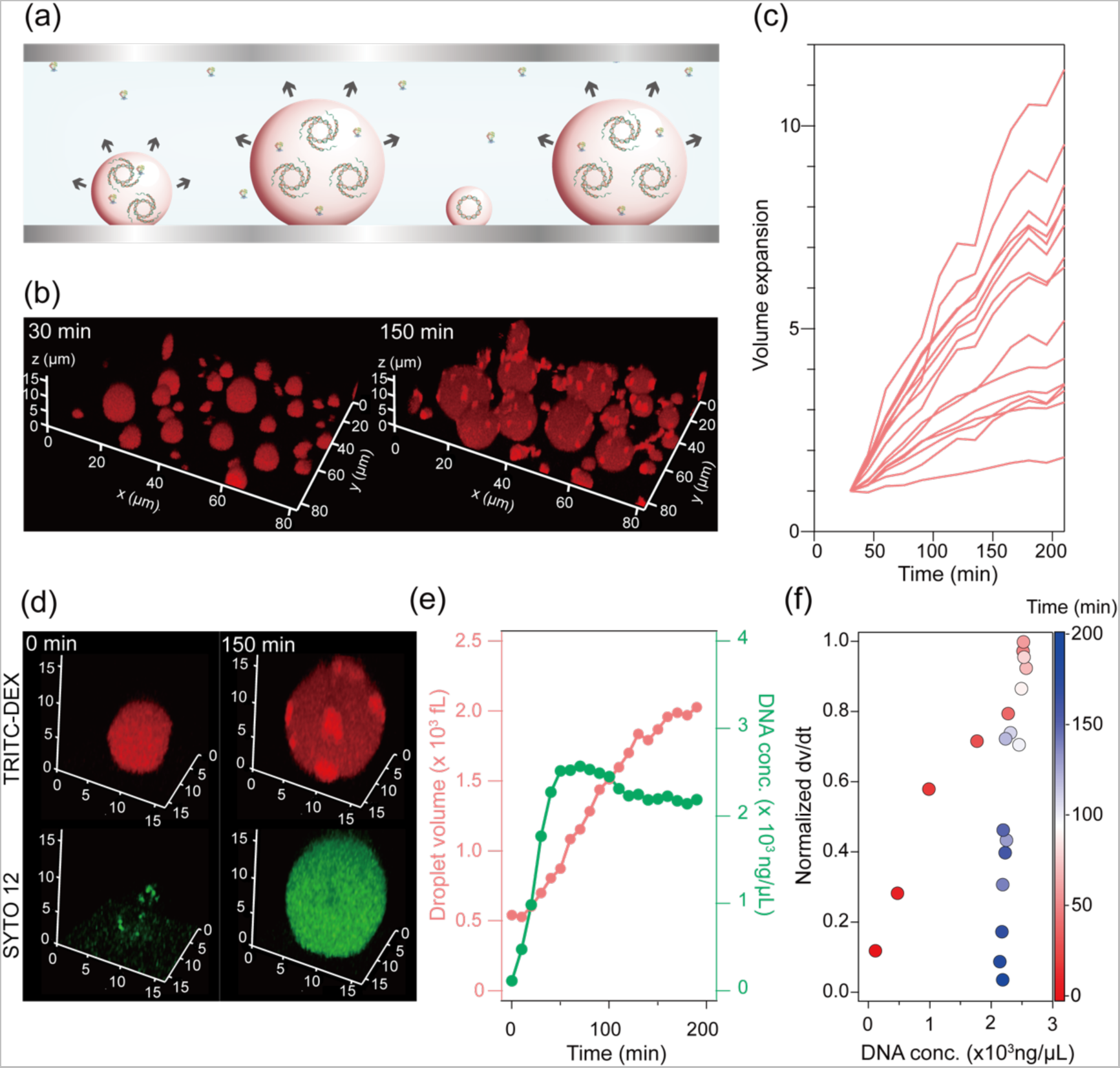
Self-growth induced by RCA. (a) Schematic image of DEX droplets self-growth induced by DNA amplification. DNA is amplified by RCA catalyzed by Equi*ϕ*29DNAP. (b) Confocal images of the protocell stained with TRITC-DEX (red). Fluorescence images show the reaction at 30min (left) and at 150min (right). (c) Time-courses of the volume of DEX droplets. The first 30 min was omitted because small droplets occasionally coalesced to other droplets in the first 30 min. The volume was normalized with that at 30 min and plotted. (d) Magnified images of DEX droplets stained with TRITC-DEX (red) and with SYTO 12 (green) at 0min (left) and at 150min (right). (e) Time-course of the volume changes and DNA concentration of the droplet shown in (d). Time-courses of other droplets are shown in Supplemental Fig. 7. (f) The normalized growth rate versus DNA concentration analyzed from the time-course in (e). The color of data points represents the incubation time, as shown in the color bar. Analyses for other droplets are also shown in Supplemental Fig. 7.

We also monitored DNA concentration inside the droplets by staining with an intercalator, SYTO 12 (Fig. 4d and Supplemental Video 3 and Supplemental Fig. 6). Fig. 4e shows representative time courses of [DNA] and the droplet volume. While [DNA] increased along with the droplet growth, [DNA] reached a plateau level even earlier than droplet growth. This trend was observed for other droplets (Supplemental Fig. 7). Interestingly, the plateau levels of [DNA] were found around 2000 ng/μL. This suggests that 2000 ng/μL is close to the upper limit for DEX droplets to accommodate DNA inside under the present condition. When the rate of volume expansion was plotted against [DNA] inside the droplets, it turned out that the growth rate monotonically increased with [DNA] until [DNA] reached the plateaus level (Fig. 4f). Considering that the rate of DNA amplification should increase in proportional to [DNA] in the early phase, it is reasonable to observe that the droplet growth is highly dependent on the internal [DNA] when it is below the upper limit.

We also tested whether RNA was also able to induce phase separation in the PEG/DEX system. When RNA with 1.75 × 10^3^ nt in length at 623 ng/µL was spiked to PEG/DEX mix at (*2.0*, *2.0*), the PEG/DEX mix underwent phase separation, generating DEX droplets (Extended Data Fig. 1a,1b). We also confirmed that *in vitro* transcription in miscible PEG/DEX mix induces phase separation (Extended Data Fig. 1c,1d). Thus, it was confirmed that *in situ* RNA production can also induce the phase separation of PEG/DEX solution as well as DNA amplification.

To seek more autonomy of the DEX droplet as the artificial cell model, cell-free transcription-translation system (TXTL) was added with the template DNA encoding Equi*ϕ*29DNAP to implement TXTL-coupled RCA system in the DEX droplets (Fig. 5a) (40). We prepared PEG/DEX ATPS mix containing the cell-free TXTL system from purified components, PURE system (41). While most of PURE system components such as ribosome, transcription factors are intrinsically enriched in the DEX-rich phase (37), Equi*ϕ*29DNAP is almost evenly partitioned in both DEX-rich and PEG-rich phases. In order to retain expressed Equi*ϕ*29DNAP inside the droplet, it was tagged with dextran binding domain (DBD) by genetic fusion (42). The growth activity of the DEX droplets was investigated in time-lapse imaging with confocal microscope. Many of droplets showed continuous growth over 720 min (Fig. 5b-d and Supplemental Video 4). The growth rate was slower than that of droplets carrying purified RCA components, probably due to the cascade reaction of TXTL-coupled RCA. Many of droplets increased the volume over 3-fold during 12 hr incubation, and some were significantly active in the growth, expanding the volume over 8-fold. When dNTPs were not added, droplets did not show obvious growth, confirming that the active growth was driven by TXTL-coupled RCA reaction (Supplemental Fig. 9).

**Fig. 5.**
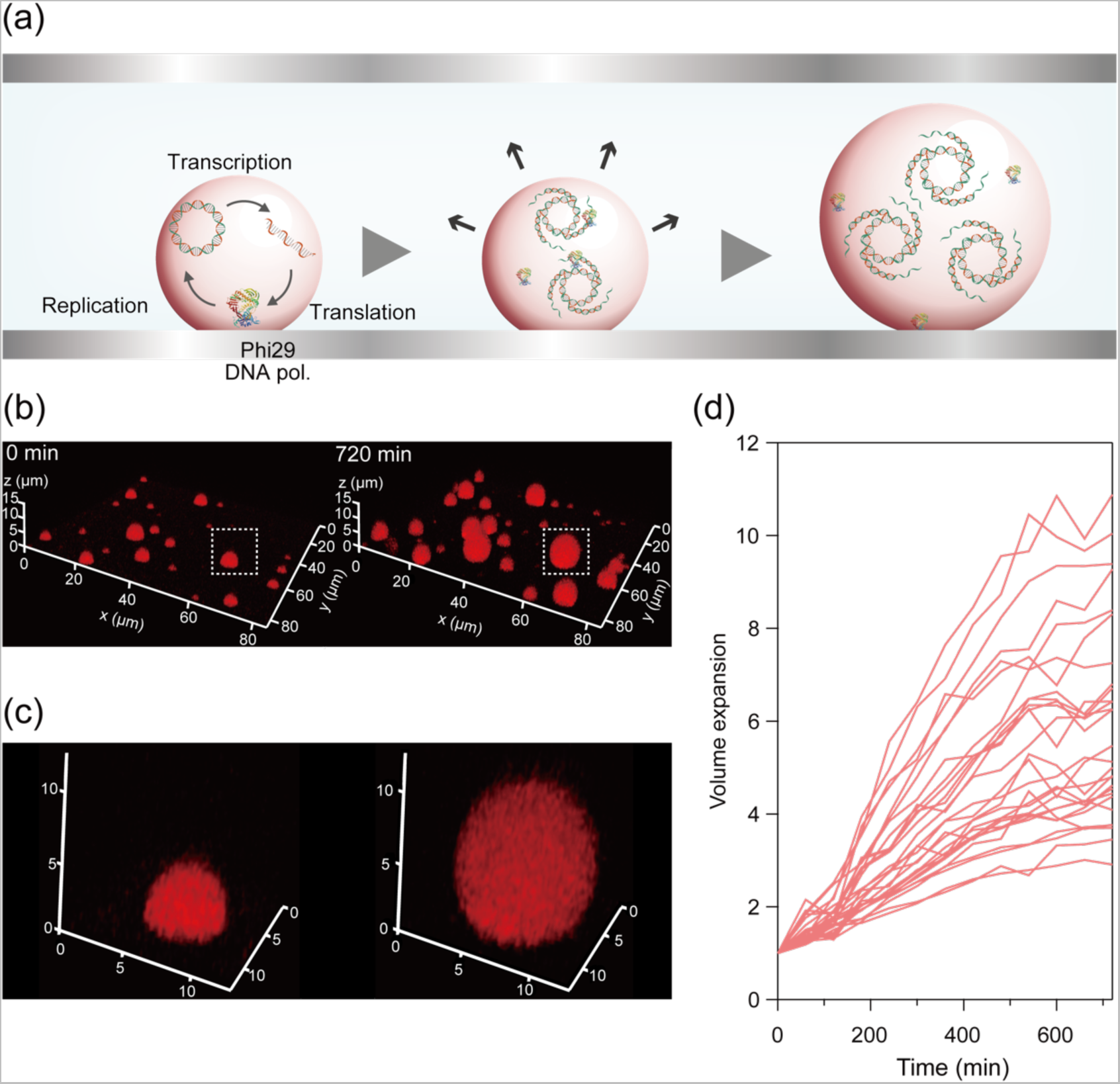
Self-growth by TXTL-coupled RCA. (a) Experimental diagram. Cell-free TXTL system (PURE system) was implemented to DEX droplet with DNA encoding Equi*ϕ*29DNAP tagged with dextran binding domain (Equi*ϕ*29DNAP-DBD). Upon the expression of Equi*ϕ*29DNAP-DBD and subsequent DNA amplification, the protocells showed the volume expansion. (b) Confocal images of the protocell stained with TRITC-DEX. Fluorescence images show the reaction at the start (0 min, left) and at 720min (right). (c) Magnified droplet image surrounded by white dashed square in (b). (d) Time-courses of self-growth of the protocells. The volume was normalized with that at 0 min.

Then, we attempted to implement a recursive DNA replication system that produce the exact same copies of template DNA in terms of length and topology. Of isothermal and recursive DNA replication systems (12,43,44), we employed the reconstituted *ϕ*29 phage genome replication system, terminal protein primed DNA amplification (TPPDA) (43), considering the simplicity and the application for a protocell model (45). TPPDA is principally composed of four components: terminal protein (TP), DNA polymerase (*ϕ*29DNAP), ssDNA binding protein (SSB), dsDNA binding protein (DSB). Prior to the growth experiment under microscope, we confirmed that TXTL-coupled TPPDA system retains the functionality to replicate DNA in PEG/DEX ATPS (Supplemental Fig. 10). To implement TXTL-coupled TPPDA system in the DEX droplet, 13 kbp linear DNA encoding genes of TP and *ϕ*29DNAP was added as the template DNA, while SSB and DSB were added as purified proteins with PURE system (Extended Data Fig. 2a). The DEX droplets implemented with TXTL-coupled TPPDA showed continuous self-growth activity over 280 min and exhibited volume expansion: 1.8 ± 0.2-fold (Extended Data Fig. 2b-d and Supplemental Video 5). Thus, the DEX droplet-based protocell model showed self-growth activity to increase the volume almost two times upon the recursive DNA amplification. The growth activity is moderate in comparison with the growth activity by TXTL-coupled RCA. This is attributable to the complex nature of TPPDA; this recursive replication system requires four protein components while RCA requires only polymerase, and thereby TPPDA should be more susceptive to the leakage effect and the interaction with PEG/DEX. In particular, we noticed that TP-attached DNA products is prone to form aggregation (Supplemental Fig. 13), that would at least partially explain the lower activity of TPPDA-driven self-growth.

Finally, we sought the possibility of self-growth activity starting from a single molecule of template DNA, aiming to approach an autonomous protocell model with evolvability that generally require a single set of gene-coding molecule. Preliminary experiments showed that RCA and TPPDA are not suitable because of the non-specific amplification of RCA and low replication activity of TPPDA. We then tested the *E. coli* genome replicasome system termed RCR system (12) that is the recursive DNA replication system composed of 25 protein components including *E. coli* DNA polymerase. We found RCR system is functional in the DEX droplet array on femtoliter reactor array device (FRAD) (42) where a million of DEX droplets are displayed on the reactor device and covered by PEG-rich phase solution (Fig. 6a), whereas RCR was not sufficiently active in free DEX droplets for unknown reasons. The template DNA was highly diluted to ensure DNA molecules were stochastically partitioned into DEX droplet reactors at a single molecule level. Along with DNA molecules, the purified components of RCR were also introduced into DEX droplet reactors. Probably because the initial DNA concentration in DEX droplet reactor was too low (0.33 ng/µL), the obvious volume expansion of DEX droplet reactors was not observed after 4 hr incubation while we observed the increment of intercalator fluorescence in some reactors. Therefore, we attempted further cultivation of the reactors by refreshing the PEG-rich solution containing RCR components for the 2nd incubation. As expected, while empty reactors retained their original volume without the volume expansion, the reactors that showed distinct fluorescence signal of DNA intercalator in the 1st incubation showed obvious volume expansion of DEX droplet, protruding upwardly (Fig. 6b,6c and Supplemental Video 6). The self-growth always accompanied with the expansion of intercalator fluorescence image, showing the self-growth was driven by internal DNA replication. The growth ratio was high, ranging from 2 to over 3 (Fig. 6d). This high growth ratio is probably due to high activity of long DNA replication by RCR. Although this system has not yet achieved the coupling with TXTL, the result suggests the possibility of autonomous growth induced by TXTL-coupled DNA replication from a single molecule of DNA.

**Fig. 6.**
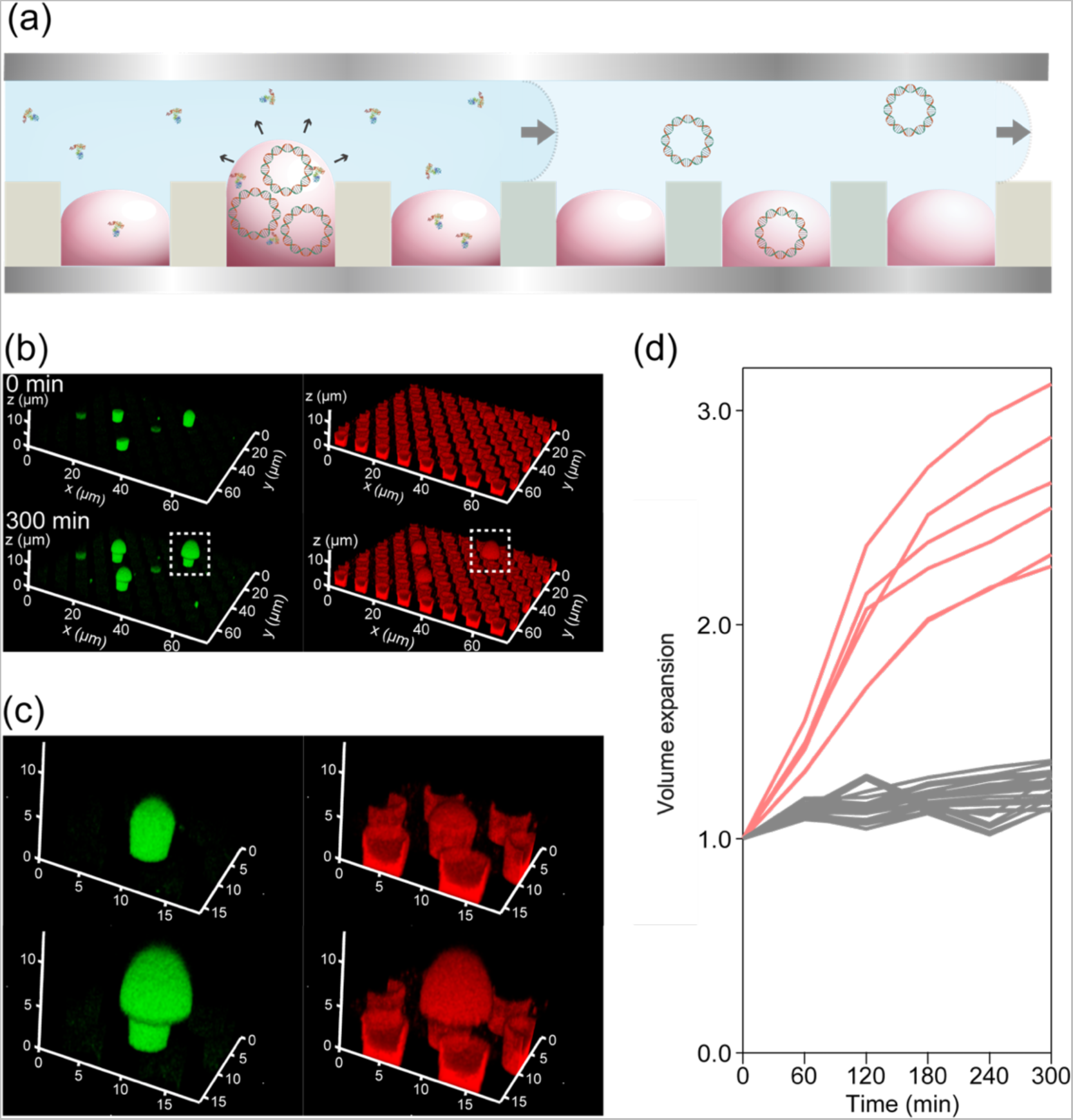
Self-growth induced by RCR DNA amplification on FRAD. (a) Experimental diagram. DEX droplets were formed on femtoliter reactor array devices (FRAD). The DNA concentration was limited to 0.7 pM, (= 0.02 DNA molecules per reactor) that ensures digital assay conditions wherein most reactors are empty, while a few reactors entrap single molecules of DNA. After the 1st round of RCR, the protocells were incubated with a fresh PEG-rich solution containing RCR components at 33 ℃ (for details, see Methods). (b) Sequential fluorescence images of protocells on FRAD during 2nd round of RCR. DNA and DEX were stained with SYTO 13 (green) and TRITC-DEX (red) at 0 min (top) and at 300 min (bottom). (c) Enlarged view of the droplet surrounded by the white dashed square in (b). (d) Time-course of self-growth of protocells. The volume was normalized with that at 0 min. Red lines represent the time-courses of the protocell with DNA, and gray ones show that of DNA-free protocells.

## Discussion

The PEG/DEX system is one of the best-characterized aqueous phase separation systems. Preferential partitioning of biopolymers, such as DNA, RNA, proteins, and protein filaments, in the DEX-rich phase has been reported (32–36). However, the reverse effect, that is, phase stabilization by such biopolymers, has not been reported. This study showed that DNA stabilize the ATPS of the PEG/DEX system and quantitively analyzed the impact of DNA on the phase separation. We also found RNA induces phase separation although the quantitative analysis was not allowed due to the difficulty of the preparation of homogeneous RNA sample at large scale. The stabilization effect was observed in narrow conditions; (*2.09, 2.09*) - (*1.77, 1.77*) for 205 kbp DNA when the titration experiment started from (*2.30, 2.30*). In addition, a large quantity of DNA or RNA (600-900 ng/µL) was required to observe the stabilization of ATPS. These might be the reason why the stabilization effect of the DNA/RNA reverse effect has not been found in previous works.

Longer DNA stabilized ATPS more efficiently. This is in accordance with the trend of the partitioning ratio in the DEX-rich phase, suggesting that stabilization and partitioning are based on a common mechanism. DEX and PEG are chemically inert polymers without charged or aromatic rings unlike droplet-forming biopolymers. Therefore, the interplay between DNA and the DEX-rich phase would be based on physicochemical principles rather than chemically specific interactions. Yoshikawa et al. have proposed that filamentous polymers/assemblies, such as double-stranded DNA and actin filaments, are more readily accommodated in larger void interspaces in the DEX-rich phase than in the PEG-rich phase (34). On the other hand, the present study found that RNA, a much more flexible polymer is also preferentially enriched in the DEX phase, implying that other mechanisms also exist.

By exploiting the phase stabilization effect of DNA, we attempted to develop self-growing protocell models. For this purpose, we implemented internal DNA/RNA synthesis reactions within DEX droplets. When RCA system was implemented, the DEX droplets showed significant growth of the droplet size, expanding the volume by 2-10 times. Simultaneous observation of the DNA amplification and droplet growth showed that the growth rate raised with the internal [DNA] and it reached the maximum when [DNA] reached the plateau level around 2000 ng/μL. These observations showed that the internal DNA concentration acts as the internal pressure to drive the droplet growth.

In pursuit of a more autonomous protocell model, we attempted to implement the ability to self-produce key enzymes for gene replication. At first, we implemented cell-free TXTL-coupled RCA. Upon the gene expression, the template DNA was amplified by the *in situ* produced Equi*ϕ*29DNAP that resultantly induced the active growth, achieving the growth ratio by 3-10 times. Then, we implemented cell-free TXTL-coupled recursive DNA replication system, TPPDA into DEX droplet. The DEX-droplets showed active growth over 280 min, expanding 1.8 times from the initial volume. Thus, we achieved the development of an autonomous protocell model that undergoes self-growth driven by internal amplification/replication of polymerase-encoding DNA coupled with gene expression.

There are other reports on self-growth activity of several protocell models. Liposome growth induced by *in situ* synthesis of lipid molecule catalyzed by chemical catalysts were actively studied (14,46). It was also attempted to build self-growing liposome systems with cell-free TXTL system to produce enzymes for lipid synthesis (14,17). Although this strategy is the straightforward for build-a-cell, the growth ratio is still modest and not coupled with gene replication. The growth of coacervate droplets was also reported in several studies in which the growth were induced by the addition or enzymatic production of inducer chemicals such as ATP or UDP (30). The implementation of gene replication systems to induce self-growth of coacervate is still challenge. Dex-droplet-based protocell models represent a completely new model with large potential to create protocells with autonomy; because the driving force for self-growing is based on simple mechanism, the enzymes for the self-growing are genetically encodable and the genetic materials such as DNA and RNA are readily retained inside of the cells. Considering these points, the present study opens a new venue to build protocell models with high autonomy and functionality.

However, it should also be mentioned that there are many critical challenges for DEX-droplet based protocell models. One major obstacle is how to implement the capability for self-division. Although the surface tension of DEX droplet is lower than that of water-in-oil droplet (34), it is still challenging to implement a mechanism to induce division of DEX droplets. In this regard, a implicative work was reported by Takinoue group (47). They have created oligo DNA droplet composed of modestly associating oligo DNA strands and succeeded in splitting droplets into two parts by introducing a restriction enzyme. Although the implementation of gene expression and gene replication are remaining challenges for their strategy, but this work is very suggestive for the consideration of the division mechanism for DEX droplet-based protocell. Symmetry-breaking structures in DEX droplets would be a key factor.

Another challenge is to gain evolvability. For evolution experiments, it is required to encapsulate DNA molecules as few as possible in each DEX droplet. Ultimately, it is desirable to partition template DNA under digital conditions (8,48), i.e. with zero or one molecule of DNA in each cell. To achieve this, it is required to create protocells that can grow even starting from a single molecule of DNA. In this regard, we have not achieved obvious growth by TXTL-coupled TPPDA from a single-molecule DNA level. To seek the possibility of self-growth activity from a single copy of DNA, we implemented RCR system in DEX droplet array displayed on FRAD, and introduced single molecules of template DNA for RCR. DEX droplets carrying a single molecule DNA showed remarkable growth activity, supporting the possibility to develop protocell model with self-growth activity. Thus, the DEX droplet-based protocell with TXTL-coupled RCR is one of the next challenges.

The results of this study also provide valuable insights into how protocells could have arisen in environments prior to the emergence of primitive cells. While the protocell model used in this study is based on polymers that did not exist in prebiotic conditions, the self-growth mechanism discovered here highlights the reverse reaction of preferential partitioning of solute molecules that, in turn, induce and stabilize phase separation. This could be a phenomenon universally observed in other phase-separating systems, including those that might have existed on primitive Earth. The self-growth mechanism demonstrated in this study is much simpler and more easily coupled with gene replication compared to liposome-based protocells. The simplicity of the self-renewal mechanism in DEX droplet-based protocells may provide an important clue to the scenario of the emergence of primitive life. However, it is crucial to examine other phase-separated systems to further understand the potential mechanisms that could have driven the formation and evolution of early protocells.

## Methods

The detailed methods are provided in supplemental information.

### Analysis of DNA partition ratio

The indicated mixtures of DNA (20 ng/µL), DEX, and PEG were prepared in 1.5 mL tubes and centrifuged at 15,000 rpm for 3 min. The mixture separated into two layers: an upper layer rich in PEG and a lower layer rich in DEX. The DNA concentrations in each phase were determined from the fluorescence signal of SYBR Gold using calibration curves, each prepared for 10 bp, 100 bp, 4 kbp, or 205 kbp DNA in 4 w/w% DEX or 4 w/w% PEG (Supplementary Fig. 1).

### Titrimetry of PEG/DEX ATPS with or without DNA

ATPS mixtures of the PEG/DEX system were prepared at the following concentrations: (*1.2, 2.5*), (*2.3, 2.3*), (*3.2, 2.0*), (*4.8, 1.5*), (*6.0, 1.2*), or (*7.0, 0.9*) with 4 kbp DNA (950 ng/μL), with 205 kbp DNA (950 ng/μL), or without DNA. 100 μL of PEG/DEX mix was serially diluted by addition of 2 μL of pure water. Each time after the dilution, the samples were centrifuged at 15,000 rpm for 3 min for the segregation of PEG-rich layer on the top and DEX-rich layer at the bottom of tubes. To enhance the visual clarity of the phase separation, DEX- and PEG-rich phases were stained with 0.1% w/w of TRITC-DEX and 0.1% w/w of FITC-PEG.

### Phase separation induced by RCA

Rolling circle amplification (RCA) was conducted in PEG/DEX mix (*2.00, 2.00*) with a commercially available kit of Equi*ϕ*29DNAP in the presence of 8 pM circular 205 kbp DNA, 0.05% TRITC-DEX, 1 µM SYTO 13, and 6N random primer. The mixture was incubated at 33 °C for 16 h in a flow cell.

### Self-growth induced by RCA

The DEX droplets were prepared by mixing of DEX-rich phase solution from (*5.0, 5.0*) and PEG-rich phase solution from (*2.5, 2.5*) containing buffer for Equi*ϕ*29DNAP at 1:50 ratio to suppress the volume ratio of DEX droplets to PEG-rich phase and thus, to avoid coalescence of DEX droplets for time-lapse imaging. The template 4.2 kbp plasmid DNA and Equi*ϕ*29DNAP were added to the DEX droplets just before infusing the solution into a flow cell assembled from two CYTOP-coated cover slips. The sample was incubated at 33℃.

### Phase separation induced by spiked RNA

A plasmid DNA, pRSET B, encoding a derivative of mScarlet protein and eight Pepper sequences (8Pepper) downstream under T7 promoter, was used as the template. RNA transcript with 1750 nt long was synthesized with a ScriptMAX^®^ Thermo T7 Transcription Kit and purified with NucleoSpin RNA Clean-up XS. The purified RNA was added at 623 ng/µL to the miscible PEG/DEX mixture (*2.00, 2.00*) with 0.05 w/w% TRITC-DEX and 20 mM HBC.

### Phase separation induced by in situ RNA transcription

A plasmid DNA, pRSET B, encoding a derivative of mScarlet protein and 8Pepper downstream under T7 promoter, was used as the template. The complete RNA transcript was approximately 1750 nt in long. 600 ng/µL of the template DNA and ScriptMAX^®^ Thermo T7 Transcription Kit was added to miscible PEG/DEX mixture (*2.00, 2.00*) with 0.05 w/w% and HBC (20 mM) and incubated at 33 °C for 16 h.

### Self-growth by TXTL-coupled RCA

The 5.2 kbp plasmid DNA encoding Equi*ϕ*29DNAP tagged with DEX-binding domain (DBD) at C-terminus fused was used as template (42). The DEX droplets were formed by mixing DEX-rich solution and PEG-rich solution prepared from (*5.0, 5.0*) or (*2.5, 2.5*) containing components for the TXTL-coupled RCA at 1:30 ratio. The template DNA and solution B for TXTL kit were added to DEX droplets just before infusing the solution into a flow cell assembled from two CYTOP-coated cover slips. The sample was incubated at 30℃.

### Self-growth by TXTL-coupled TPPDA DNA replication

The 13kbp linear DNA encoding *ϕ* 29DNAP, TP with *ϕ* 29 origins was used as a template (45). The DEX droplets were formed by mixing DEX-rich solution and PEG-rich solution prepared from (*8.0, 8.0*) or (*3.0, 3.0*) containing the components for TXTL-coupled TPPDA at 1:30 ratio. The template DNA, purified SSB, purified DSB, and solution B for TXTL kit were added to the DEX droplets just before infusing the solution into a flow cell assembled from two CYTOP-coated cover slips. The sample was incubated at 30℃.

### Self-growth induced by RCR DNA amplification on FRAD

RCR buffer, RCR enzyme mix were provided from Moderna Enzymatics Co., Ltd. The flow cell was assembled from FRAD and a non-fabricated SiO2 glass coated with CYTOP separated by approximately 80 μm double-sided tape. First, DEX solution (3% (w/w) DEX, 0.01% (w/w) TRITC-DEX) was introduced into the flow cell. Then, PEG solution (3% (w/w) PEG equilibrated with DEX, RCR buffer) with 12 kbp DNA encoding OriC sequence was injected in the flow cell to flush out extra dextran solution and form DEX droplets in each chamber. Subsequently, PEG solution (3% (w/w) PEG equilibrated with DEX, RCR buffer) with RCR enzyme mix was introduced into the flow cell, and the solution on the device was incubated at 33℃ for 4 hr (1st round of RCR). After that, a fresh PEG solution (3% (w/w) PEG equilibrated with DEX, RCR buffer) with RCR components and 1 μM SYTO13 was introduced into the flow cell and incubated at 33℃ (2nd round of RCR). The protocells were observed under confocal microscope during the incubation.

### Microscopic imaging and image analysis

See supplemental information.

## Acknowledgements

We thank Ryoko Yaginuma for technical supports. We also thank Yuki Goto Kyoto University and Kazuhito V. Tabata from The University of Tokyo for generously providing the HBC reagent. This work was financially supported by CREST, Japan (JPMJCR19S4 to H. N. and JPMJCR18S6 to M. S.) from Japan Science and Technology Agency (JST), GteX program (JPMJGX23B1 to H. N.) from JST, Grant-in-Aids for Scientific Research (S) (JP19H05624 to H. N.) from Japan Society for the Promotion of Science (JSPS) and Grant-in-Aid for Research Fellows (22J22681 to M. Y.) from JSPS.

## Author contributions

H. N. conceived the idea of the research and supervised it. Y. M. designed the experiments. Y. M. and M. Y. conducted the experiments and analyzed the data. M. S. prepared the materials for RCR and provided technical supports for the RCR experiment. H. N., Y. M, and M. Y. wrote the manuscript.

---

**Extended Data Fig. 1.**
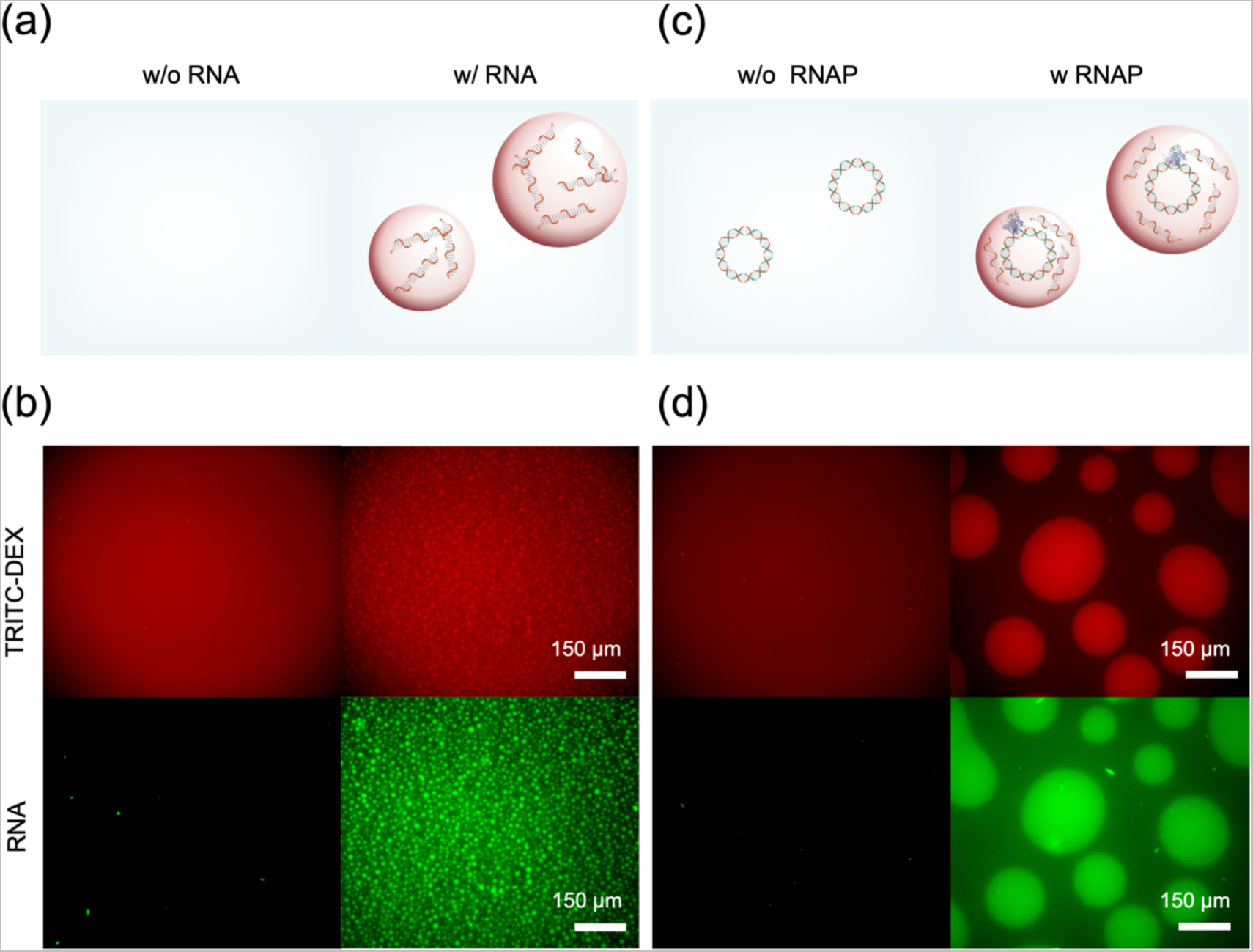
Phase separation induced by RNA. (a) Experimental diagram of (b). (b) Phase separation induced by spiked RNA. Purified RNA was not added (left) or added to (*2.0, 2.0*) PEG/DEX system at 623 ng/µL. Fluorescence images were obtained by staining RNA containing Pepper sequence with HBC (green) and DEX with TRITC-DEX (red). (c) Experimental diagram of (d). (d) Phase separation induced by *in situ* RNA transcription. Miscible PEG/DEX mix at (*2.0, 2.0*) containing 600 ng/µL template DNA were incubated with *in vitro* transcription system without (left) or with RNA polymerase (right). Fluorescence images were obtained by staining RNA containing Pepper sequence with HBC (green) and DEX with TRITC-DEX (red).

**Extended Data Fig. 2.**
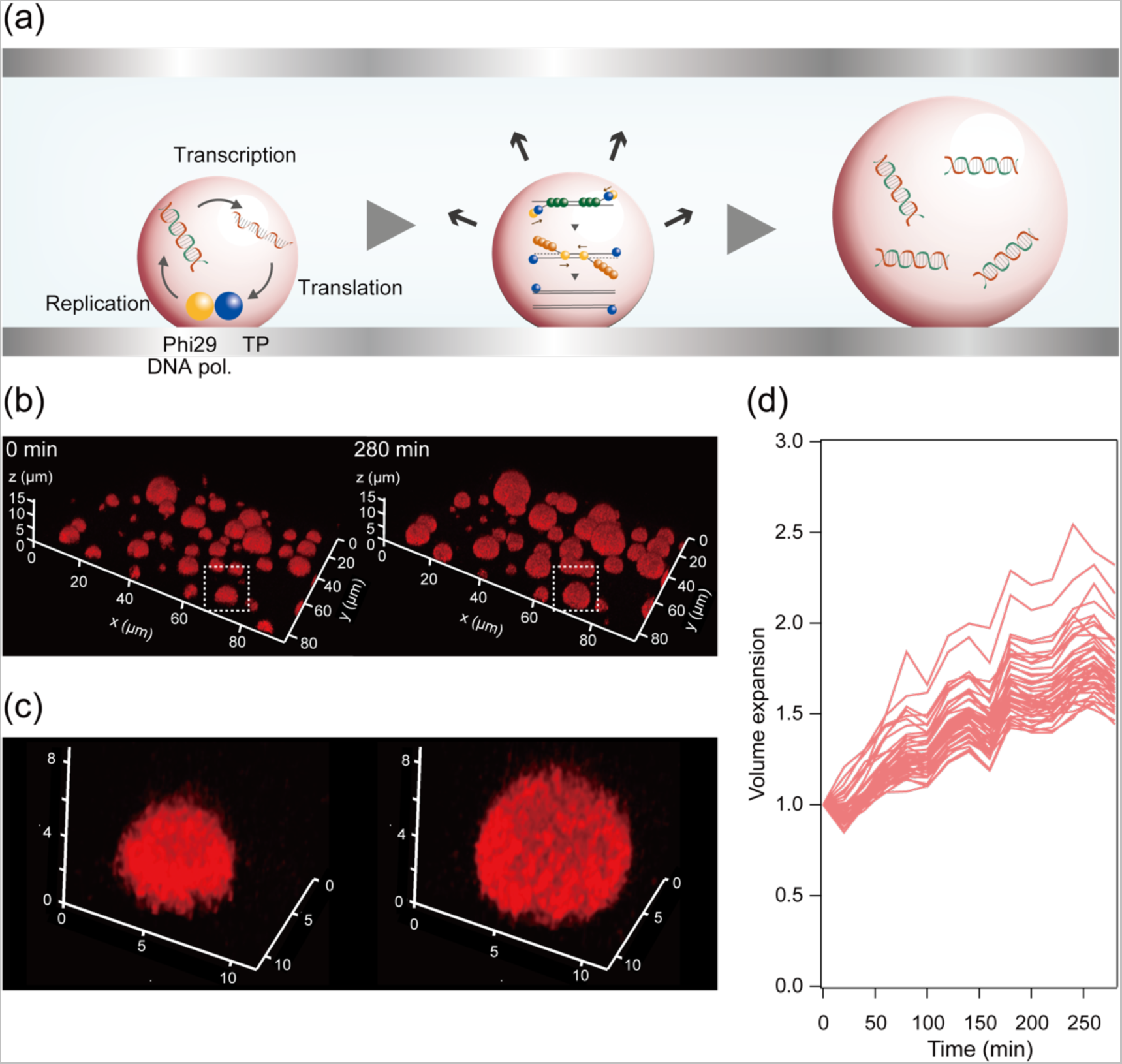
Self-growth by TXTL-coupled TPPDA. (a) Experimental diagram. Protocells containing TXTL system and terminal protein primed DNA amplification (TPPDA) system were incubated. The TXTL-coupled TPPDA system in this study was composed of template DNA encoding terminal protein and *ϕ*29DNAP, and purified proteins: ssDNA binding protein (SSB) and dsDNA binding protein (DSB). (b) Confocal images of the protocell stained with TRITC-DEX. Fluorescence images show the reaction at the start (0 min, left) and at 280min (right). (c) Magnified images of DEX droplets surround by dashed square border in (b) at 0 min (left) and at 280min (right). (d) The time-course of self-growth of protocells. The volume was normalized to that at 0 min.

