## Supplemental Information for "Self-growing protocell models in aqueous two-phase system induced by internal DNA replication reaction"

#### Table of contents

1. Extended Data Figure legends

2. Supplemental Video Legends

3. Supplemental information of materials and methods

4. Supplemental Figures

5. References

### 1. Extended Data Figure legends

#### Extended Data Fig. 1 Phase separation induced by RNA transcription

(a) Experimental diagram of (b). (b) Phase separation induced by spiked RNA. Purified RNA was not added (left) or added to (*2.0, 2.0*) PEG/DEX system at 623 ng/μL. Fluorescence images were obtained by staining RNA containing Pepper sequence with HBC (green) and DEX with TRITC-DEX (red). (c) Experimental diagram of (d). (d) Phase separation induced by in situ RNA transcription. Miscible PEG/DEX mix at (*2.0, 2.0*) containing 600 ng/μL template DNA were incubated with in vitro transcription system without (left) or with RNA polymerase (right). Fluorescence images were obtained by staining RNA containing Pepper sequence with HBC (green) and DEX with TRITC-DEX (red).

#### Extended Data Fig. 2 Self-growth by TXTL-coupled TPPDA

(a) Experimental diagram. Protocells containing TXTL system and terminal protein primed DNA amplification (TPPDA) system were incubated. The TXTL-coupled TPPDA system in this study was composed of template DNA encoding terminal protein and $\phi$29DNAP, and purified proteins: ssDNA binding protein (SSB) and dsDNA binding protein (DSB). (b) Confocal images of the protocell stained with TRITC-DEX. Fluorescence images show the reaction at the start (0 min, left) and at 280min (right). (c) Magnified images of DEX droplets surround by dashed square border in (b) at 0 min (left) and at 280min (right). (d) The time-course of self-growth of protocells. The volume was normalized to that at 0 min.

#### 2. Supplemental video legends

##### Supplemental video 1.

Epi-fluorescence microscopy video showing the formation of DEX-rich droplets from miscible PEG/DEX solutions. Fluorescence images were obtained by staining DNA and DEX with SYTO 13 (green) and TRITC-DEX (red), respectively. Total duration of recording is 780 min.

##### Supplemental video 2.

Confocal microscopy video showing the active growth of DEX-rich droplets induced by RCA reaction. Fluorescence images were obtained by staining DEX with TRITC-DEX (red). Total duration of recording is 210 min.

##### Supplemental video 3.

Confocal microscopy video showing the active growth of DEX-rich droplets induced by RCA reaction. Fluorescence images were obtained by staining DNA and DEX with SYTO 12 (green) and TRITC-DEX (red), respectively. Total duration of recording is 190 min.

##### Supplemental video 4.

Confocal microscopy video showing the active growth of DEX-rich droplets induced by transcription-translation coupled RCA reaction. Here, the plasmid DNA encoding phi29 DNA polymerase tagged with Dextran Binding Domain was used as the template. Fluorescence images were obtained by staining DEX with TRITC-DEX (red). Total duration of recording is 720 min.

##### Supplemental video 5.

Confocal microscopy video showing the active growth of DEX-rich droplets induced by transcription-translation coupled TPPDA (terminal protein primed DNA amplification) reaction. Here, the 13kbp linear DNA encoding phi29 DNA polymerase and terminal protein was used as the template. Fluorescence images were obtained by staining DEX with FITC-DEX (red). Total duration of recording is 280 min.

##### Supplemental video 6.

Confocal microscopy video showing self-growth of DEX droplet induced by RCR in FRAD. Here, 12 kbp cicular DNA encoding OriC was encapsulated in DEX droplet under digital conditions. At the start of the video (0 hour), the sample had already undergone 4 hour RCR reaction at 33℃. Total duration of recording is 300 min.

#### 3. Supplemental information of materials and methods

##### Supplemental information of materials

DEX (MW: 450,000–650,000 Da) from *Leuconostoc* spp., tetramethylrhodamine isothiocyanate-dextran (TRITC-DEX; MW: 500,000 Da), and PEG (MW: 35,000 Da) were purchased from Sigma-Aldrich, St. Louis, MI, USA; FITC-methyl poly(ethylene glycol) (FITC-PEG; MW: 30,000 Da), CreativePEGWorks, USA; CYTOP, Asahi-glass, Japan; SYBR Gold, Green Fluorescent Nucleic Acid Stain (SYTO13), EquiPhi29™ DNA polymerase, EquiPhi29™ DNA polymerase Reaction Buffer (10X), and Exo-Resistant Random Primer, Thermo Fisher Scientific, US; and ScriptMAX® Thermo T7 Transcription Kit, Toyobo, Japan; PUREExpress In Vitro Protein Synthesis Kit, Ribonucleotide Solution Set, NEW ENGLAND Biolabs, USA; 6N random primer, FASMAC, Japan; dNTPs Mixture, Nippon Gene, Japan, NucleoSpin RNA Clean-up XS, Takara-bio, Japan. The proteins and chemicals used for RCR were prepared as previously reported (1). The DNA for the titration experiment was prepared as follows. DNA fragments of 10 bp and 100 bp long were prepared by annealing complementary oligonucleotide pairs; 4 kbp, digesting a plasmid DNA encoding *E. coli* alkaline phosphatase in pRSET B (Thermo Fisher Scientific) (2); 12 kbp, generating a plasmid DNA in-house by assembling 1.2 kbp, 1.8 kbp, 1.9 kbp, and 2.3 kbp fragments, each encoding mScarlet, amp$^r$, replication origin, and *E. coli OriC* sequences, and 2.6 kbp and 2.4 kbp fragments obtained from digesting 85 kbp plasmid DNA, pOri80; 205 kbp, pMSR 227 (1) . The composition of 1X Energy Mix is as follows; 0.36 mM each 2 natural L-amino acids, 70 mM Potassium-glutamate, 0.375 mM Spermine, 25 mM Creatine-phosphate potassium salt, 0.518 g/L *E.coli* tRNA, 0.1 M HEPES-KOH pH8.0, 0.79 mM Hemi-magnesium glutamate, 6 mM Dithiothreitol (3). The composition of rNTPmix is as follows; 18.75 mM ATP, 12.5 mM GTP, 6.25 mM UTP/CTP. The composition of RCR mixture is previously reported (1).

#### Microscopic imaging

Microscopic images were taken with a TCS SP8 confocal laser scanning microscopy with ×100 objective lens (Leica Microsystems, Germany) equipped with a white light laser (Leica Microsystems, Germany) except Fig.3 and Extended Data Fig. 1. The filter sets in each experiment are shown below.

| | Fluorescent DEX | $\lambda_{ex}$ | $\lambda_{em}$ | DNA intercalator | $\lambda_{ex}$ | $\lambda_{em}$ |
|---|---|---|---|---|---|---|
| Fig. 1 | TRITC-DEX | 555 nm | 571-625 nm | SYBR Gold | 495 nm | 500-549 nm |
| Fig. 4 | TRITC-DEX | 552 nm | 573-625 nm | SYTO 12 | 495 nm | 510-550 nm |
| Fig. 5 | TRITC-DEX | 552 nm | 573-625 nm | | | |
| Fig. 6 | TRITC-DEX | 554 nm | 570-620 nm | SYTO 13 | 488 nm | 505-545 nm |
| Extended Data Fig. 2 | FITC-DEX | 495 nm | 510-550 nm | | | |

Fig. 3 and Extended Data Fig. 1 were taken with an epi-fluorescence microscope (Olympus IX83, Olympus Co., Tokyo, Japan) equipped with an sCMOS camera (Andor Neo, Andor Technology, Belfast, Northern Ireland), an LED light source (X-Cite XYLIS, Excelitas Technologies), appropriate filter sets ($\lambda_{ex}$ = 489 nm and $\lambda_{em}$ = 508 nm for SYTO 13 and HBC, $\lambda_{ex}$ = 558 nm and $\lambda_{em}$ = 583 nm for TRITC-DEX).

#### Image analysis

The time-lapse of DEX droplets were analyzed with ImageJ. The volume of each droplet was calculated from z-stack images. DNA concentration within each droplet was estimated based on the fluorescence intensity of SYTO 12 at a height of 5 μm above the glass bottom using a calibration curve (Supplemental Fig. 6).

#### Supplemental method for self-growth induced by RCA

DEX-rich phase solution was prepared from the PEG/DEX mix (*5.0, 5.0*) containing 1x EquiPhi29$^{TM}$ DNA polymerase Reaction Buffer, 1 mM DTT and 0.05 wt% TRITC-DEX. PEG-rich phase solution was prepared from the PEG/DEX mix (*2.5, 2.5*) containing 1x EquiPhi29 DNA polymerase Reaction Buffer,1 mM DTT, 1 mM each dNTPs, 10 μM Exo-Resistant Random primer, 0.05 wt% TRITC-DEX, 20 μM SYTO 12 (only when monitoring the DNA concentration inside the DEX droplet). The sample was prepared by mixing 1 μL DEX-rich phase solution, 50 μL PEG-rich phase solution, 1 μL 1.5 ng/μL 4.2 kbp plasmid DNA, and 2 μL 1.5 μM EquiPhi29 DNA polymerase. The flow cell, assembled from two CYTOP-coated cover slips, was preincubated with 1/10 diluted solution of solutionB for TXTL kit (PURE Express, NEB) to make the bottom cover slip sticky and stably attach the droplet onto it. The solution was infused into a flow cell and incubated at 30 °C.

#### Self-growth by TXTL-coupled RCA

DEX-rich phase solution was prepared from the PEG/DEX mix (*5.0, 5.0*) containing solution A (PURE Express, NEB), 1 x Energy mix and 0.01 wt% TRITC-DEX. PEG-rich phase solution was prepared from the PEG/DEX mix (*2.5, 2.5*) containing 1x Energy mix, 1.2 mM each dNTPs, 10 μM Exo-resistant primer, solution A at a ratio of 1/25 of the total volume, rNTP mix at a ratio of 1/50 of the total volume, and 0.01 wt% TRITC-DEX. The protein synthesis solution was composed of 1 μL DEX-rich phase solution, 30 μL PEG-rich phase solution, 1 μL 60 ng/μL DNA encoding EquiPhi29 DNA polymerase fused with DBD, and 3 μL solution B. The flow cell, assembled from two CYTOP-coated cover slips, was preincubated with blocking solution containing 5 mg/ml BSA and 0.1 wt% S386. The sample solution was infused into the flow cell assembled from two CYTOP-coated cover slips and incubated at 30 °C.

#### Self-growth by TXTL-coupled TPPDA DNA replication (Extended Data Fig.2)

The 13kbp linear DNA encoding $\phi$29DNAP, TP with phi29 origins was used as a template (4). DEX-rich phase solution was prepared from the PEG/DEX mix (*8.0, 8.0*) containing 20mM ammonium sulfate, 1 x Energy mix, 0.1% FITC-DEX. PEG-rich solution was prepared from the PEG/DEX mix (*3.0, 3.0*) containing 1 x Energy mix, 20mM ammonium sulfate, 1mM dNTPs each, rNTPmix at a ratio of 1/50 of the total volume, solution A at a ratio of 1/25 of the total volume, 0.01% FITC-DEX. The sample was prepared by mixing 1 μL DEX-rich phase solution and 30 μL PEG-rich phase solution,3 μL 16.6 nM template DNA, 2.77 μL 279 μM DSB, 2.61 μL 335 μM SSB, 1.55 μL 1.6 M NaCl, and 7.75 μL solutionB. The flow cell, assembled from two CYTOP-coated cover slips, was preincubated with blocking solution containing 5 mg/ml BSA and 0.1 wt% S386. The solution was infused into a flow cell and incubated at 30 °C.

#### Preparation of DSB

With modifications, DSB was purified according to a previous report (5). His6-TEV-DSB was expressed in *E. coli* Rosetta™ 2(DE3) pLysS Singles™competent cells. Cell lysate was loaded onto a Ni-NTA column with loading buffer (50 mM Tris·HCl (pH 7.5), 7 mM β-mercaptoethanol, 0.5 mM EDTA, 5% glycerol, and 0.8 M NaCl). Elution was achieved using the elution buffer (50 mM Tris·HCl pH 7.5, 0.5 mM EDTA, 7 mM β-mercaptoethanol, 5% glycerol, 0.2 M NaCl, and 500 mM imidazole). The eluted fraction was 1/10 diluted with Buffer A (50 mM Tris·HCl, pH 7.5, 7 mM β mercaptoethanol, 1 mM EDTA, 5% glycerol) to reduce the NaCl concentration to 20 mM for loading to a Cytiva HiTrap™ Heparin HP column. Following washing the column with Buffer A, His6-TEV-DSB was eluted using Buffer A with 0.4 M NaCl. Subsequently, the elute was 1/10 diluted with Buffer A and loaded onto a Cytiva HiTrap™ Q HP column. After washing the column with Buffer A, His6-TEV-DSB was eluted using Buffer A with 0.6 M NaCl. Intact DSB was obtained by the cleavage of His6-TEV tag.

#### Preparation of SSB

With modifications, SSB was purified according to a previous report (6). SSB, fused with His6-tag, twin-strep-tag, and SUMO-tag at the N-terminus, was expressed in Rosetta™ 2(DE3)pLysS Singles™ competent cells. Cell lysate was loaded onto a streptactin column and washed with buffer A with 0.8M NaCl. Subsequently, SUMO protease solution (50 mM Tris-HCl (pH 7.5), 1 mM DTT, 1 mM EDTA, 20 nM SUMO protease, and 0.15% (w/w) NP-40) was added to the resin, and SUMO-tag was cleaved by overnight incubation at 4 °C. After the recovery of intact SSB, it was loaded onto a Cytiva HiTrap™ Heparin HP column and eluted in the flowthrough. Following this, SSB was loaded onto a Cytiva HiTrap™ Q HP column. After washing the column with Buffer A, SSB was eluted using Buffer A with 0.1 M NaCl.

#### 4. Supplemental figures

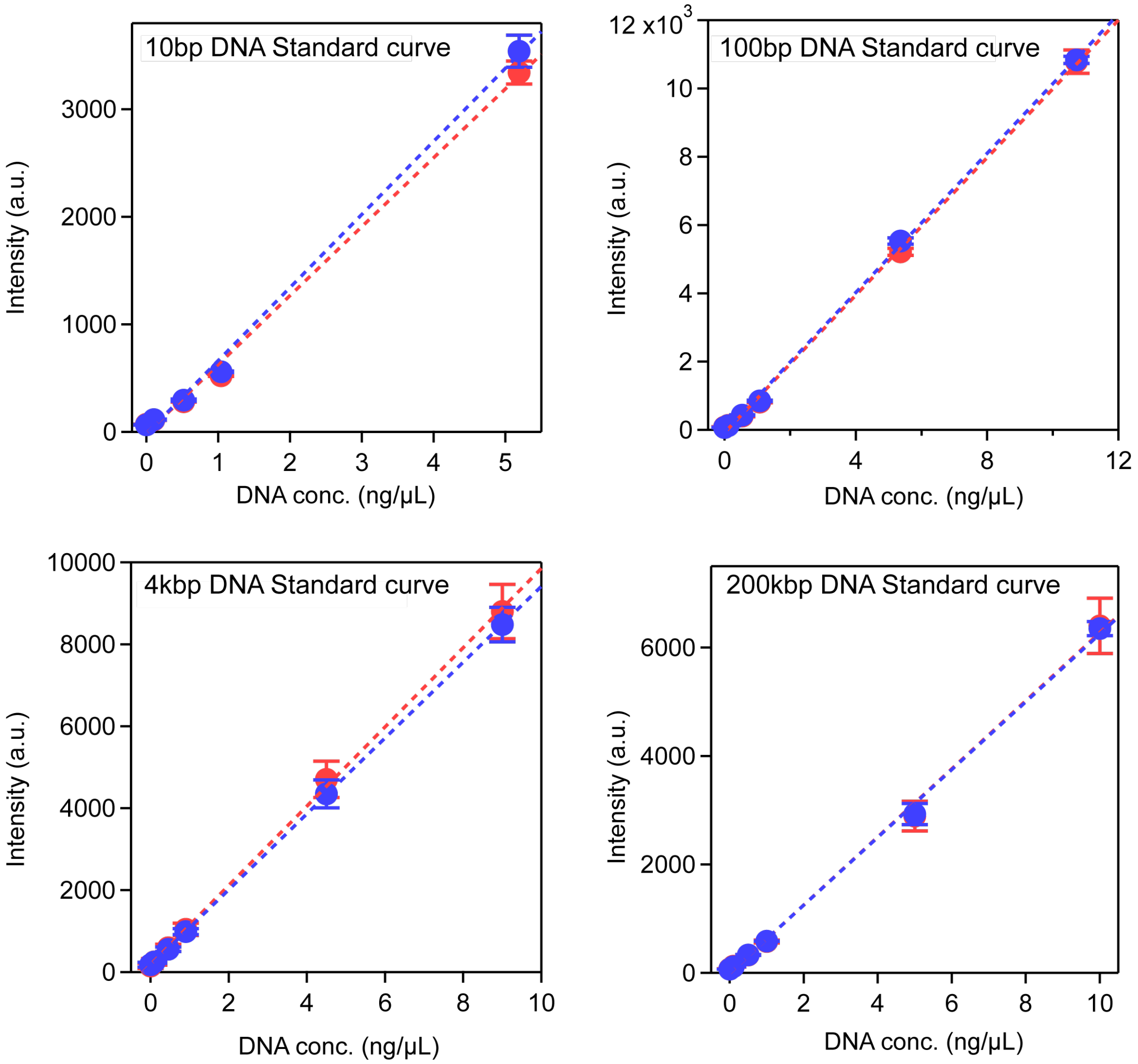


**Supplemental Fig. 1**

Calibration curves of dsDNA concentrations in 4 w/w % DEX (red) and 4 w/w % PEG (blue) using SYBR Gold for four different dsDNA lengths: 10 bp, 100 bp, 4.2 kbp, and 205 kbp. Each data point represents the mean of three independent measurements, with error bars indicating the standard deviation. The dotted lines represent the best-fit linear regression. Concentrations of DNA in Fig. 1 were quantified using these calibration curves.

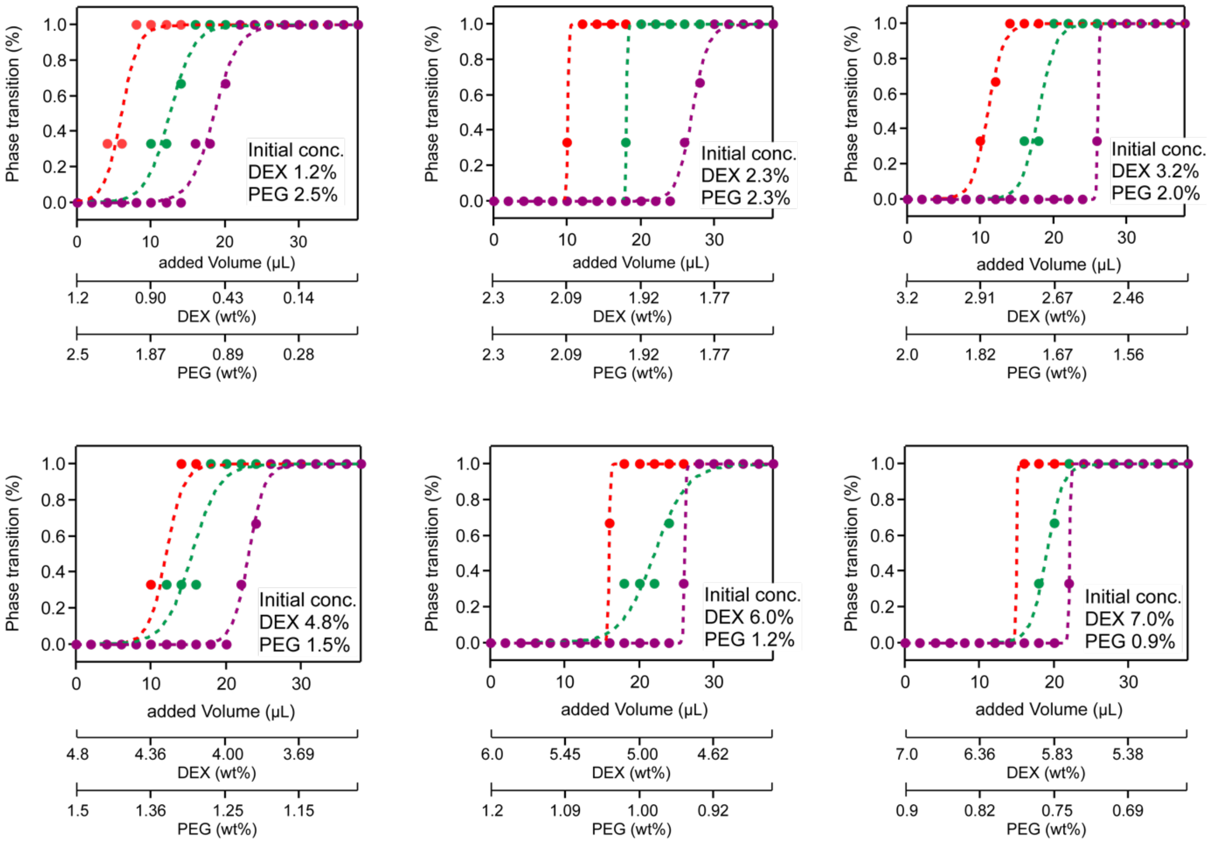


**Supplemental Fig. 2.**

Variations in phase transition probabilities at different initial concentrations of PEG/DEX, as observed in the titration experiments illustrated in Fig. 2. The red line represents the results without DNA, green for 4 kbp DNA, and purple for 205 kbp DNA.

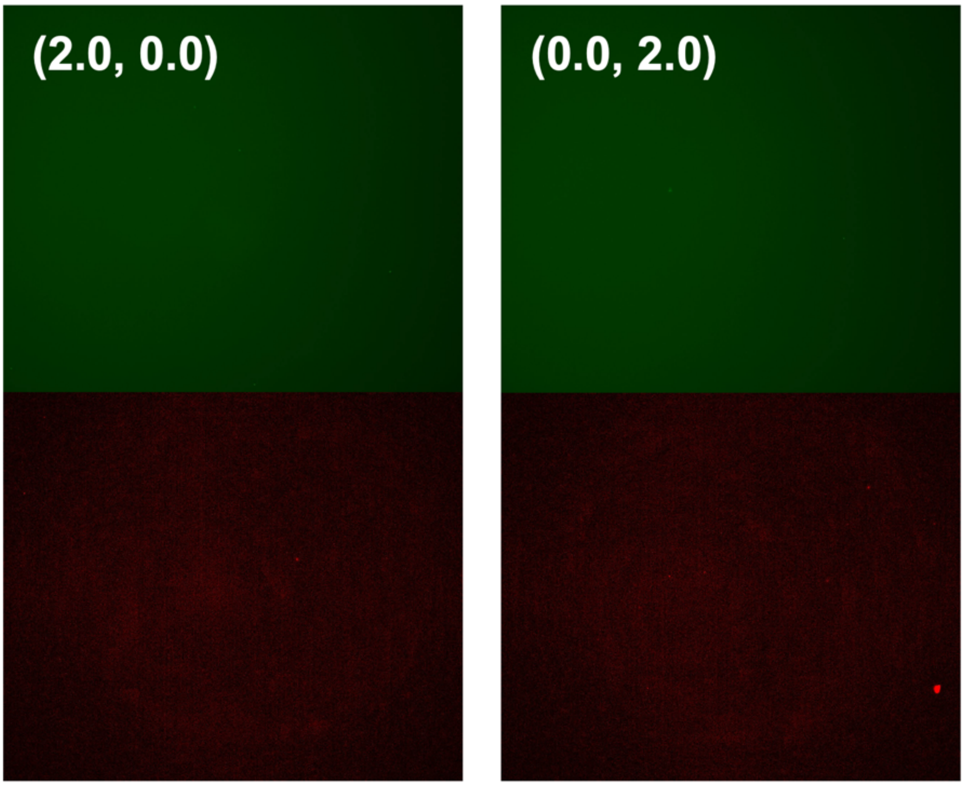


**Supplemental Fig. 3**

Fluorescence images taken after 16 hours reaction of RCA using EquiPhi29 DNA polymerase, under conditions where either DEX or PEG (but not both) was present at a concentration of 2.0%. DNA and DEX were stained with SYTO 13 (green, top) and TRITC-DEX (red, bottom), respectively.

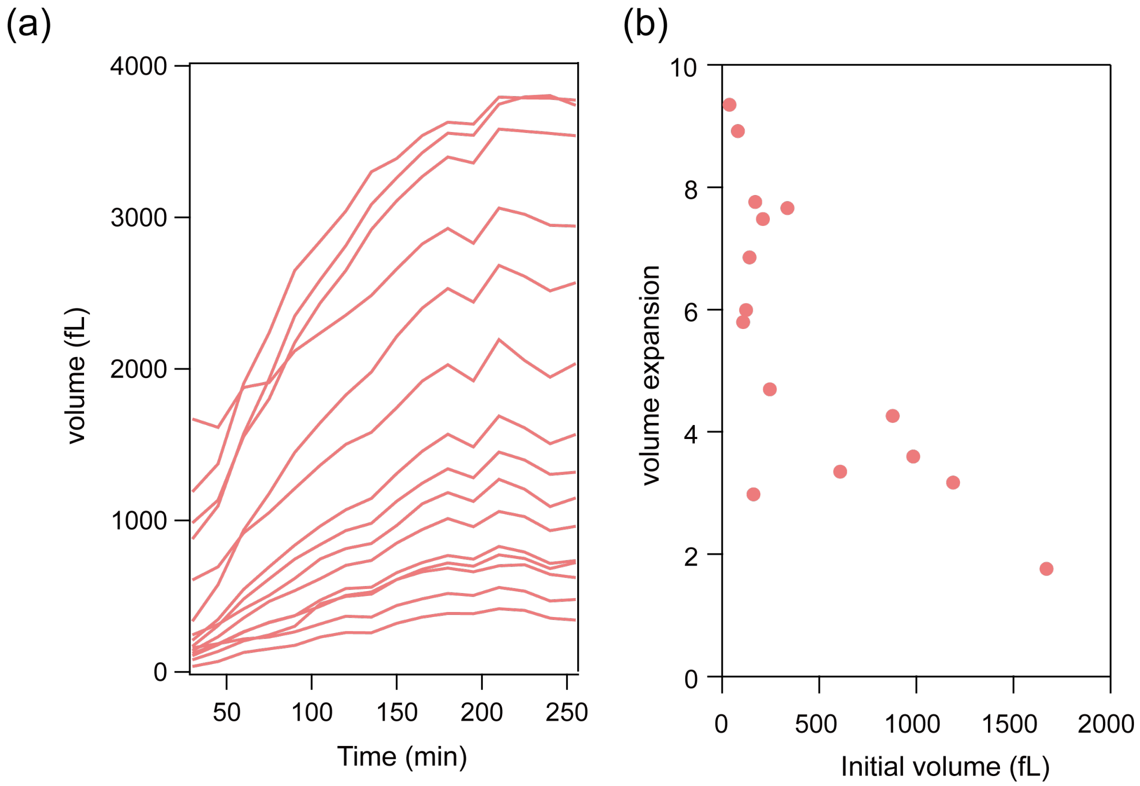


**Supplemental Fig. 4.**

(a) Time-courses of the volume of DEX droplets by RCA with Equiphi29 DNA polymerase. The first 30 minutes were omitted because small droplets occasionally coalesced with other droplets during this period. (b) The correlation between initial droplet volume and volume expansion at when droplet volumes reached plateau level. Volume expansion is calculated as the ratio of the volume at t=255 minutes to the that at t=0 minutes.

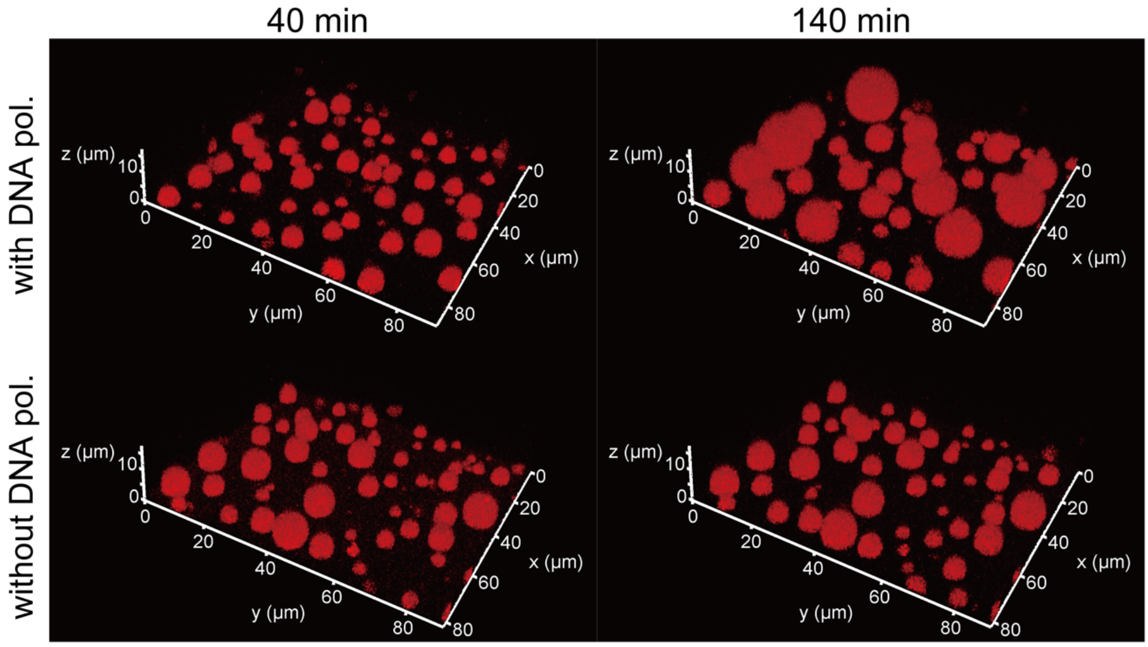


**Supplemental Fig. 5**

Representative 3D confocal fluorescence images of self-growing DEX droplets induced by RCA with (top) or without (bottom) EquiPhi29 DNA polymerase. The images show DEX droplets at 40 minutes (left) and at 280 minutes (right) of incubation. DEX is stained with TRITC-DEX (red).

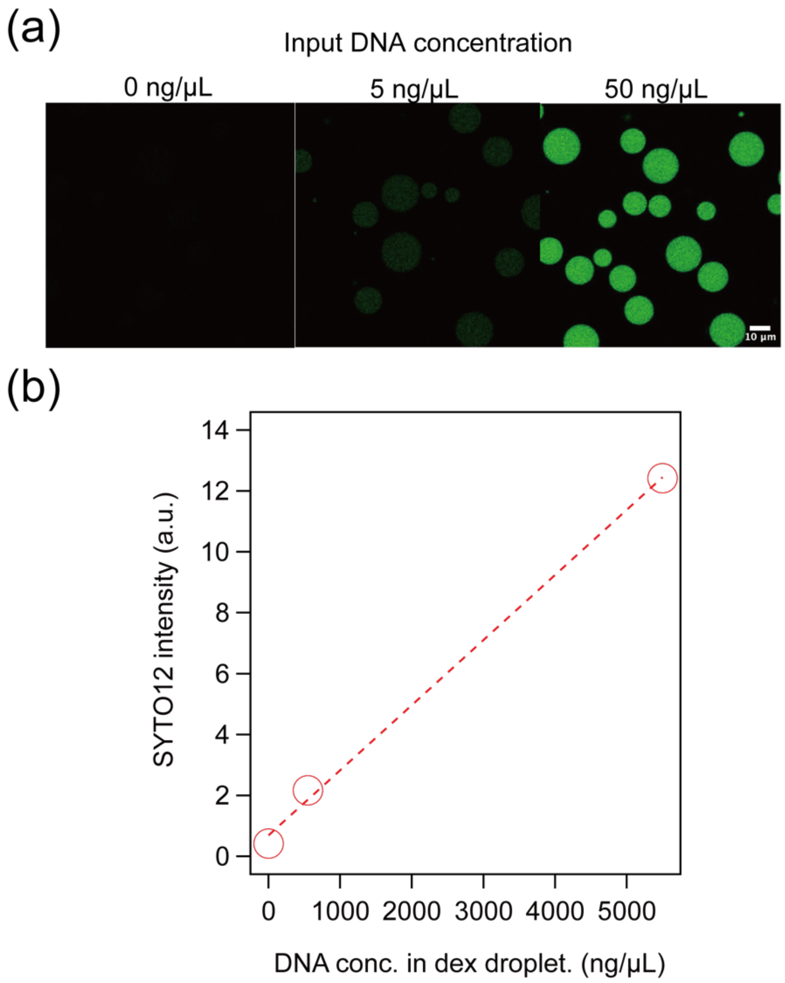


**Supplemental Fig. 6**

(a) Confocal microscopy images showing the enrichment of DNA stained with SYTO 12 (green) within DEX droplets at varying input concentrations under the experimental conditions of Fig. 4: 0 ng/μL (left), 5 ng/μL (middle), and 50 ng/μL (right). The fluorescence intensity increases with higher DNA concentrations. The scale bar represents 10 μm.

(b) Calibration curve of DNA concentration within DEX droplets from fluorescence images (Supplemental Fig. 6a). The DNA concentration inside the DEX droplet is calculated by multiplying the enrichment factor and input DNA concentration. The enrichment factor was calculated as follows: A sample with the same composition as in the microscopy images above was prepared in a tube with DNA. The DEX-rich phase and PEG-rich phase were separated, and the DNA concentration in each phase was quantified by qPCR. The DNA concentration of homogeneously mixed solution was 0.38 ± 0.03 ng/μL, that in the DEX-rich phase was 41.8 ± 1.0 ng/μL, and that in the PEG-rich phase was 0.0002 ng/μL. The enrichment factor is calculated as the ratio of DNA concentration in the DEX-rich phase to that of homogeneously mixed solution : 110.

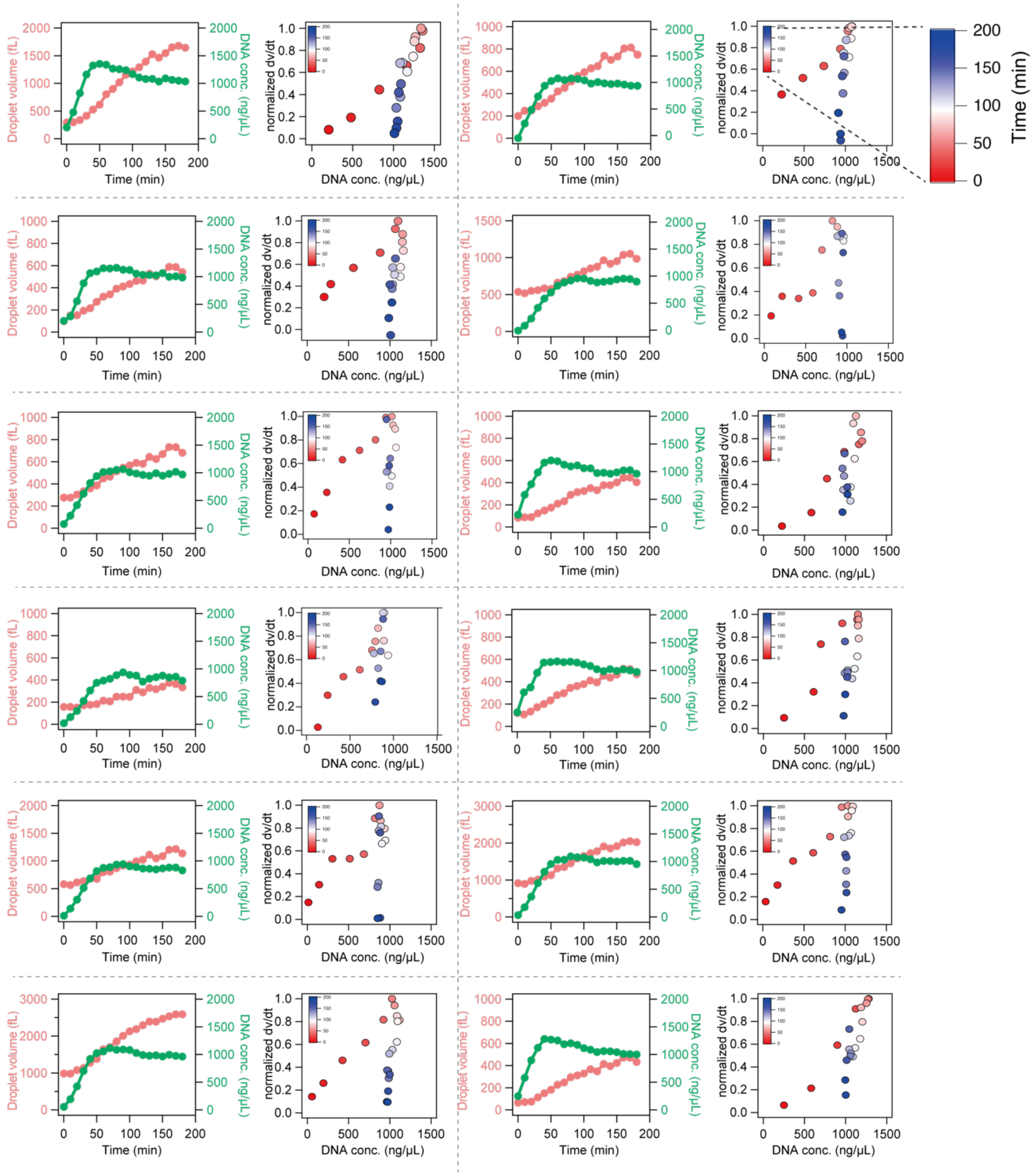


**Supplemental Fig. 7**

Supplemental figure of Fig. 4e-f. Time-courses of the volume change and DNA concentration in the droplet for other 12 droplets (left), and the normalized growth rate versus DNA concentration analyzed from the time-course (right).

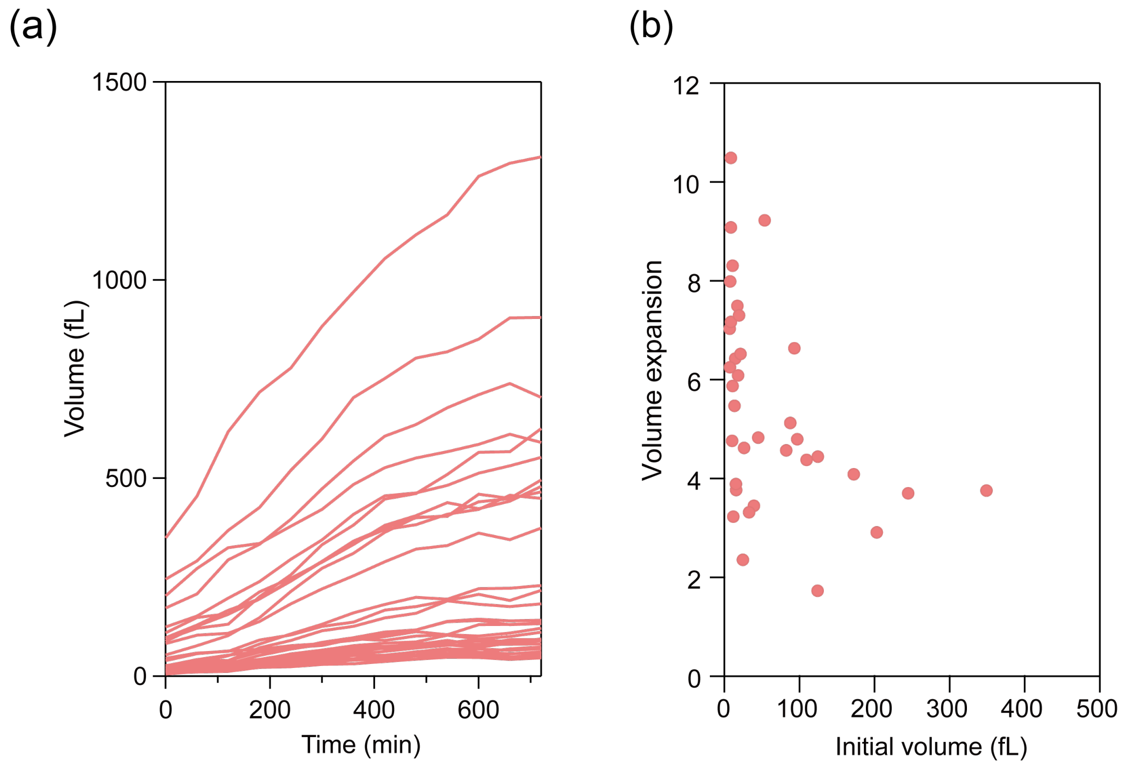


**Supplemental Fig. 8**

(a) Time-courses of the volume of DEX droplets by TXTL-coupled RCA. PURE system was implemented to DEX droplet with DNA encoding Equi$\phi$29DNAP tagged with dextran binding domain (Equi$\phi$29DNAP-DBD). Upon the expression of Equi$\phi$29DNAP-DBD and subsequent DNA amplification, the protocells showed the volume expansion.

(b) The correlation between initial droplet volume and volume expansion at when droplet volumes reached plateau level. Volume expansion is calculated as the ratio of the volume at t=720 minutes to the volume at t=0 minutes. Moderate tendency was observed that smaller droplets were more active for volume expansion as well as the DEX droplet self-growth by RCA.

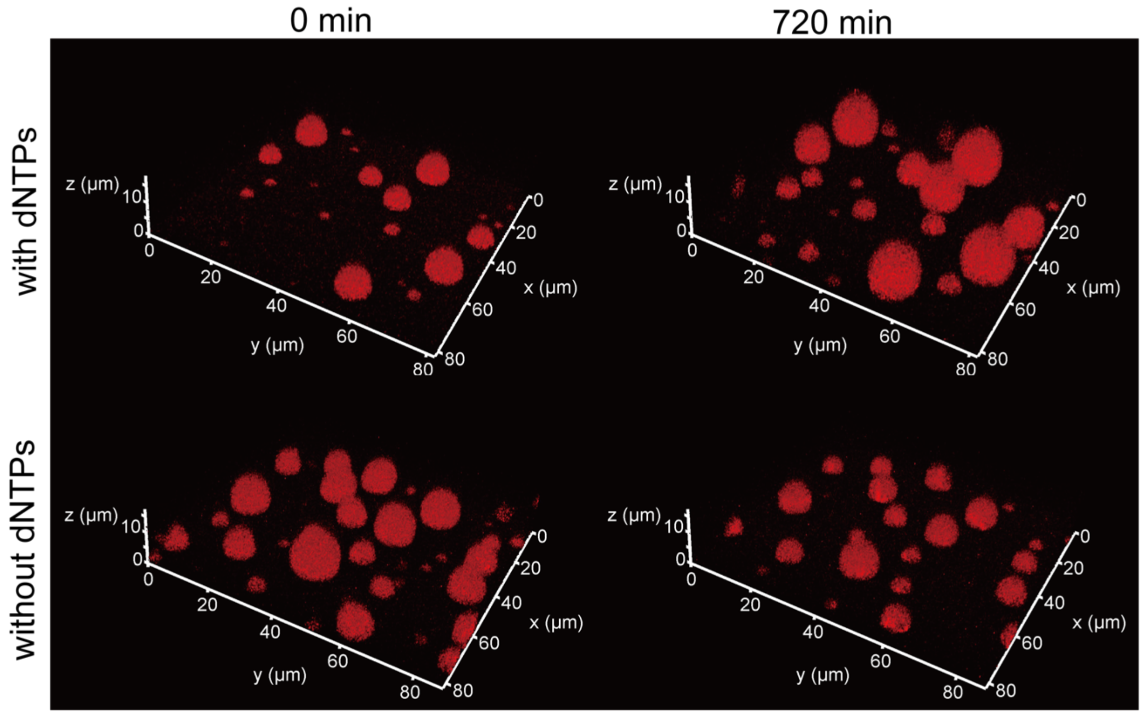


**Supplemental Fig. 9**

Representative 3D confocal fluorescence images of self-growing DEX droplets induced by TXTL couple RCA with (top) or without (bottom) dNTPs. PURE system was implemented to DEX droplet with DNA encoding Equi$\phi$29DNAP tagged with dextran binding domain (Equi$\phi$29DNAP-DBD). The images show DEX droplets at 0 minutes (left) and at 720 minutes (right) of incubation. DEX is stained with TRITC-DEX (red).

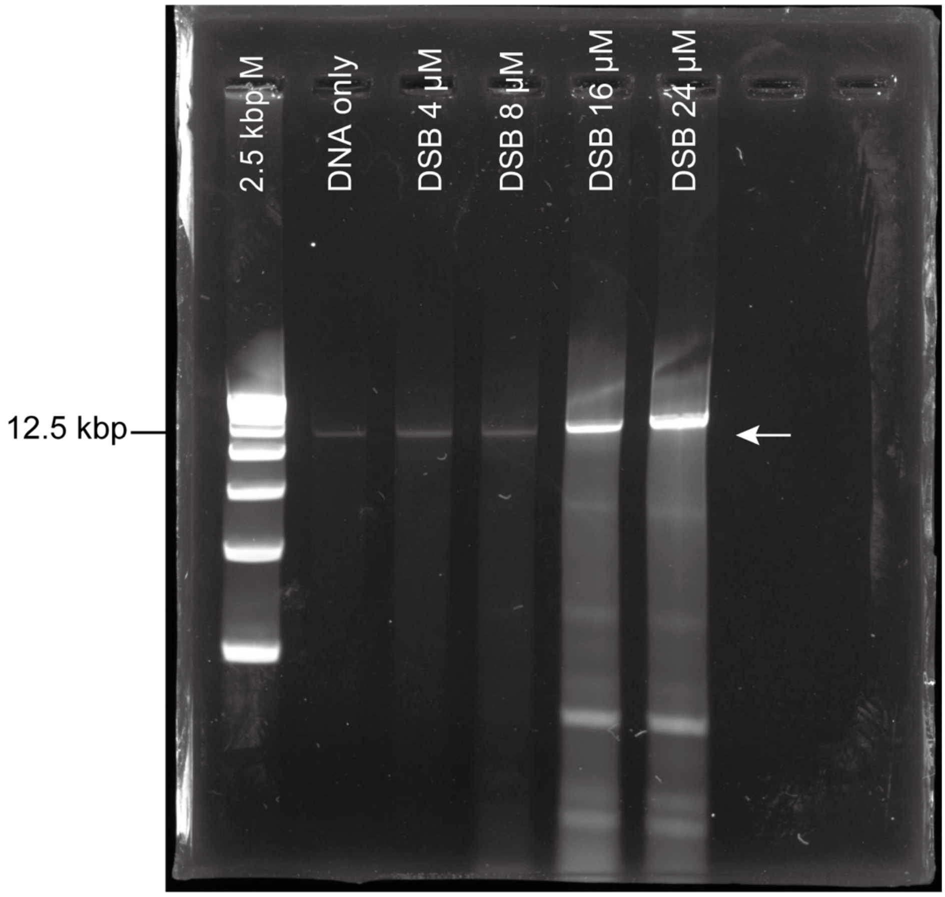


**Supplemental Fig. 10**

The TXTL-coupled TPPDA system retained the functionality to replicate DNA in PEG/DEX ATPS. Replication products in PEG/DEX ATPS were visualized on an agarose gel after RNase and Proteinase K treatments, followed by column purification. A 13 kbp linear DNA encoding the genes for TP and ϕ29 DNAP, with ϕ29 origins, was added as the template DNA to DEX droplets, while SSB and DSB were added as purified proteins with the PURE system. In each reaction, 4, 8, 16, and 24 μM of DSB were supplemented. The first lane on the gel shows the 2.5 kbp DNA marker. The second lane on the gel shows the amount of input DNA. The arrow indicates the full-length 13 kbp linear template DNA.

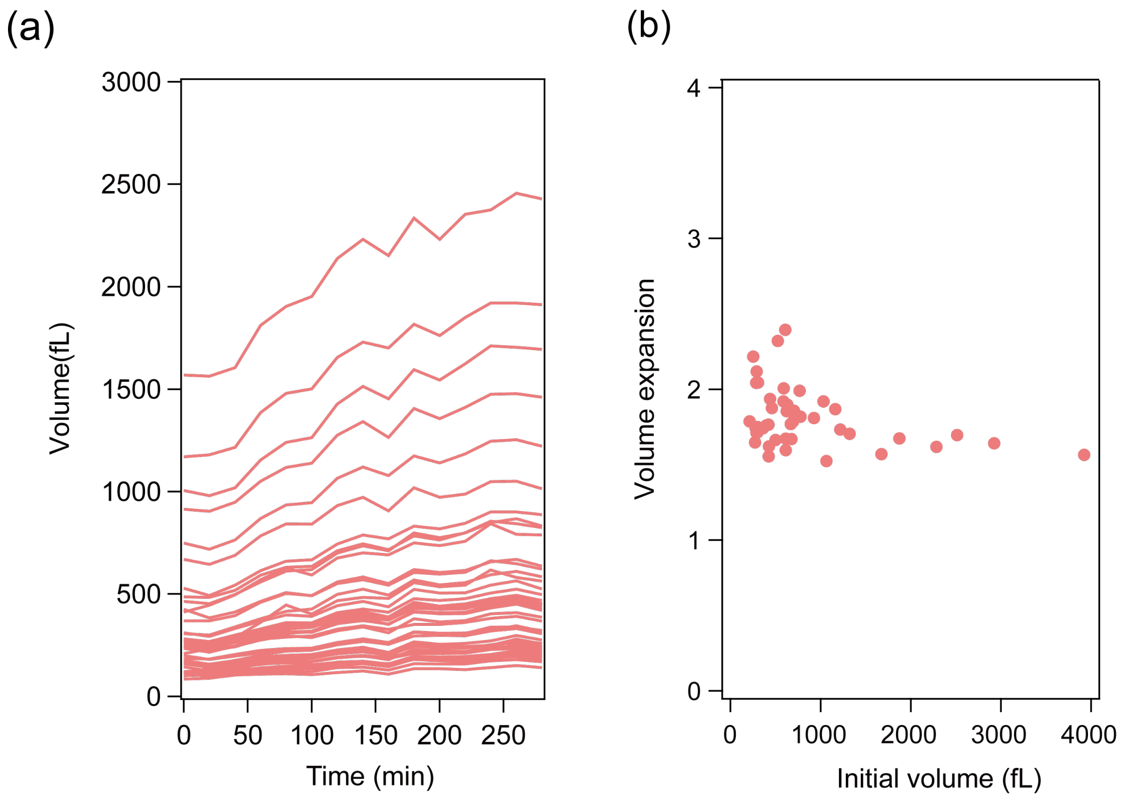


**Supplemental Fig. 11**

(a) Time-course of DEX droplet volume expansion by TXTL-coupled TPPDA reaction. Protocells containing TXTL system and terminal protein primed DNA amplification (TPPDA) system were incubated. The TXTL-coupled TPPDA system in this study was composed of template DNA encoding terminal protein and DBD-tagged $\phi$29DNAP, and purified proteins: SSB and DSB.

(b) The correlation between initial droplet volume and volume expansion at when droplet volumes reached plateau level. Volume expansion is calculated as the ratio of the volume at t=280 minutes to the volume at t=0 minutes.

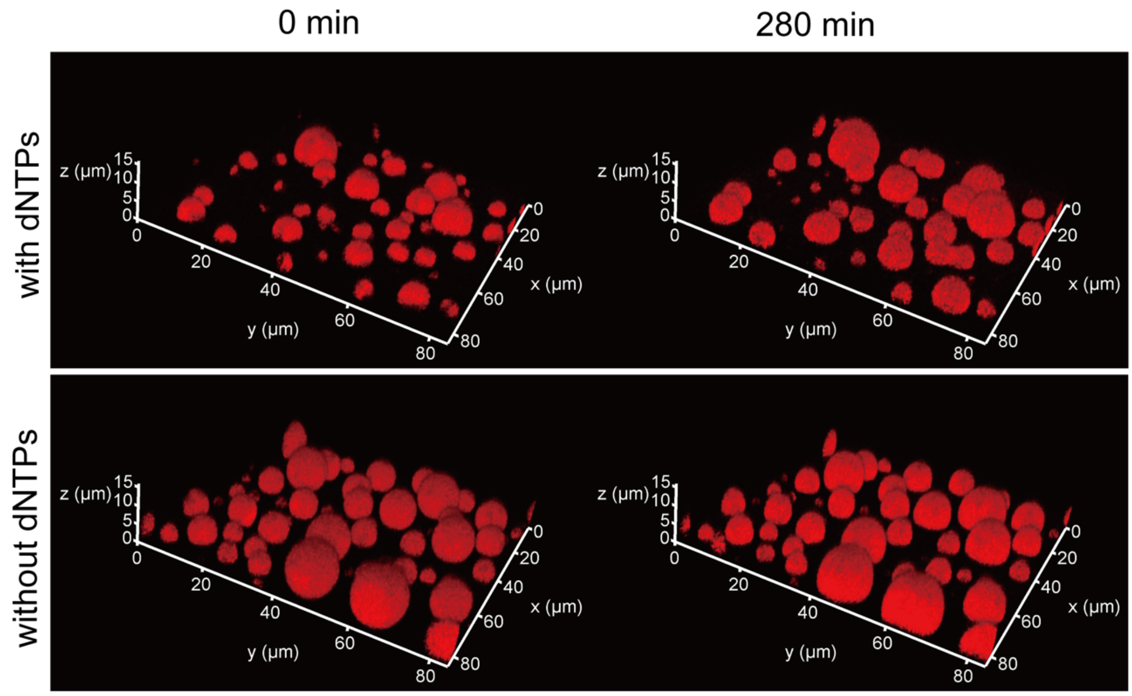


**Supplemental Fig. 12.**

Representative 3D confocal fluorescence images of self-growing DEX droplets induced by TXTL-coupled TPPDA with (top) or without (bottom) dNTPs. A 13 kbp linear DNA encoding the genes for TP and ϕ29 DNAP, with ϕ29 origins, was added as the template DNA to DEX droplets, while SSB and DSB were added as purified proteins with the PURE system. The images show DEX droplets at 0 minutes (left) and at 280 minutes (right) of incubation. DEX is stained with TRITC-DEX (red).

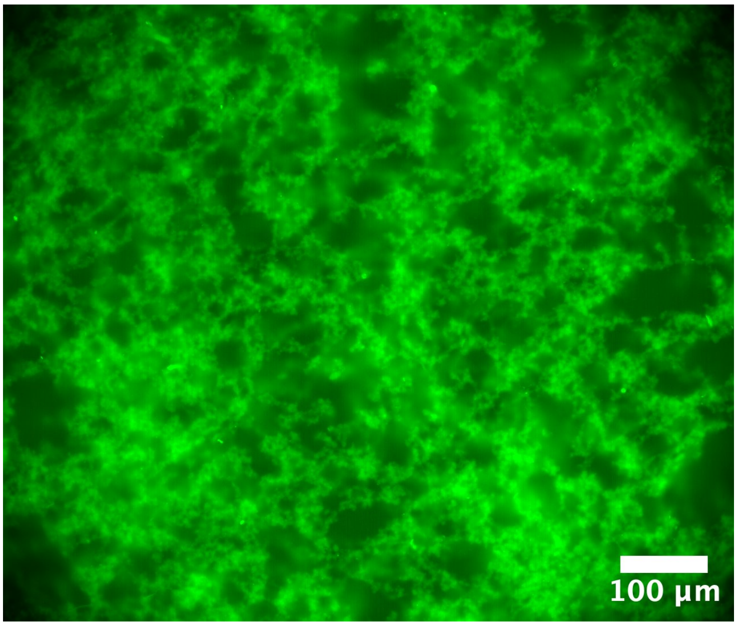


**Supplemental Fig. 13.**

Fluorescent microscopy images of TPPDA replication products after RNase treatment. The replication products were stained with SYTO 82 (green). TP-attached replication products form fiber-like aggregation.

w/ DNA (λ = 0.1)

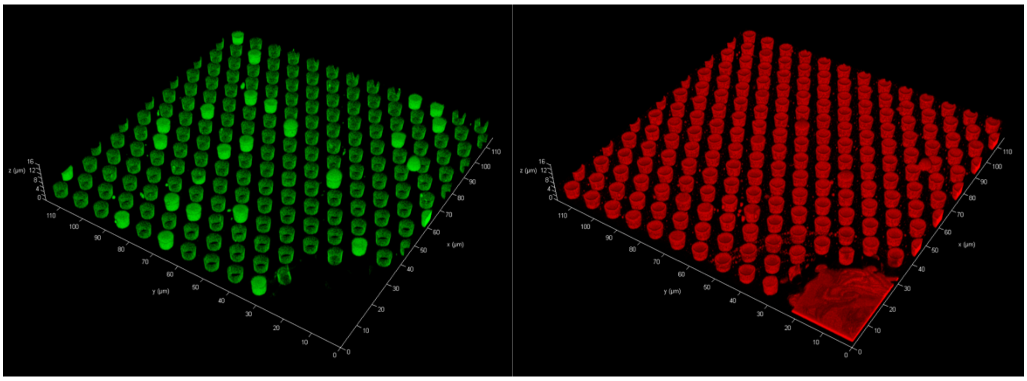

w/o DNA

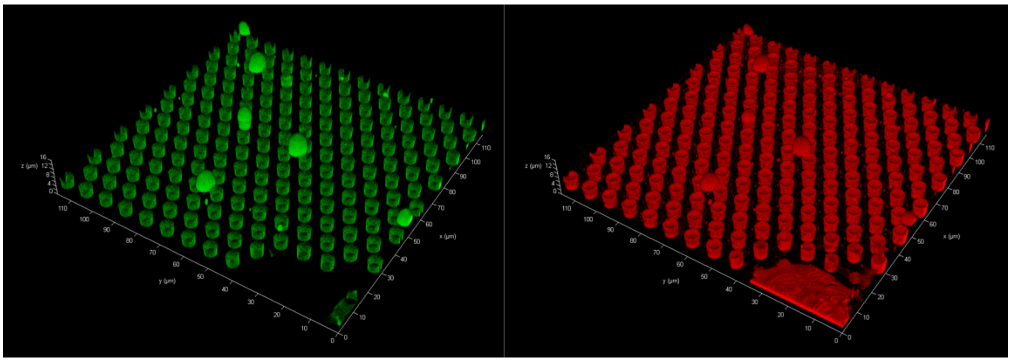

**Supplemental Fig. 14.**

Representative 3D confocal fluorescence images of self-growing DEX droplets induced by RCR with (top) or without (bottom) template DNA. The images show after 4 h incubation.
